# LC-MS profiling of *prmt-1* and *prmt-5* knockout *C. elegans* reveals PRMT-1 substrates and global proteome remodeling

**DOI:** 10.64898/2026.02.13.705869

**Authors:** Ali Basirattalab, Dylan C. Wallis, Nicolas G. Hartel, Melina Jalalifarahani, Carolyn M. Phillips, Nicholas A. Graham

## Abstract

Although protein arginine methylation regulates diverse biological processes, it remains understudied relative to other post-translational modifications. Here, we analyzed *C. elegans prmt-1* and *prmt-5* null mutants using LC-MS proteomics to map PRMT methylation substrates and to quantify the effects of PRMT knockout on global protein abundance. High-pH strong cation exchange fractionation was used to enrich methylated peptides, and parallel analysis of whole cell lysates was used to measure global protein abundance. Quantitative methyl-proteomics identified 31 PRMT-1-dependent methyl-arginine peptides from 15 proteins with several arginine residues demonstrating dramatic decrease in both monomethyl- and asymmetric dimethyl-arginine abundance. Whole-proteome profiling revealed that *prmt-1* knockout caused broad remodeling of the worm proteome with changes linked to DNA replication/cell-cycle programs, protein folding, and amino acid metabolism. Although *prmt-5* knockout affected similar biological pathways to *prmt-1* knockout, the effects on the C. elegans proteome were more modest. Together, these data connect PRMT-dependent methylation changes to proteome remodeling in a whole-animal model, support previous work suggesting that PRMT-1 is the dominant Type I PRMT in *C. elegans*, and provide a resource for studying how PRMT-1 and PRMT-5 shape protein regulation *in vivo*. All raw data have been deposited in the PRIDE database with accession number PXD074042.

## Introduction

Protein methylation is a fundamental post-translational modification (PTM) occurring on lysine and arginine residues that shapes cellular signaling, protein stability, and subcellular localization^1,2^. Lysine can accept one, two, or three methyl groups on its ε-amino side chain through the activity of lysine methyltransferases (KMTs), which use S-adenosylmethionine (SAM) as the methyl donor^3^. Protein arginine methyltransferases (PRMTs) catalyze the transfer of methyl groups from SAM to arginine residues, generating ω-N^G^-monomethyl arginine (MMA) and either asymmetric (ω-N^G^,N^G^-dimethylarginine, ADMA) or symmetric (ω-N^G^,N′^G^-dimethylarginine, SDMA) dimethylarginine marks^4^. In mammals, nine PRMT family members are classified into three types based on catalytic activity: type I enzymes (PRMT1, PRMT2, PRMT3, PRMT4/CARM1, PRMT6, PRMT8) catalyze MMA and ADMA, type II enzymes (PRMT5, PRMT9) catalyze MMA and SDMA, and the sole type III enzyme (PRMT7) produces only MMA^5^. Arginine methylation influences transcriptional regulation^6^, pre-mRNA splicing^7^, and signal transduction^8^, and other diverse cellular processes largely by modulating protein-protein and protein-nucleic acid interactions, for example through recruitment of Tudor-domain reader proteins^9^. Although many PRMT substrates contain glycine and arginine rich (GAR) motifs (RGG/RG) that are preferentially methylated^5^, substrate recognition *in vivo* is not dictated solely by this consensus sequence and can differ substantially among individual PRMTs^6^. Dysregulation of PRMT activity is implicated in numerous diseases ranging from cancer to neurological disorders^10^.

Among the PRMT family members, PRMT1 and PRMT5 account for the majority of arginine methylation in mammalian cells. PRMT1 is the dominant type I enzyme and is responsible for most ADMA modifications^11,12^, preferentially targeting RGG/RG motifs within intrinsically disordered regions of proteins including RNA-binding proteins^13^ and proteins involved in transcription, DNA repair, and signal transduction^14^. PRMT1 is highly conserved across eukaryotes from yeast to mammals, and loss of PRMT1 is lethal in mammals, underscoring its critical cellular role^15^. PRMT5 is the primary type II PRMT, catalyzing the bulk of SDMA modifications^16^. In humans and frogs (*Xenopus laevis*), PRMT5 is frequently found in a heterooctomeric complex composed of 4 PRMT5 proteins and 4 MEP50 (WDR77) proteins, an association that enhances methyltransferase activity and contributes to substrate engagement and regulation. However, this structure is not conserved across all metazoans. In *C. elegans*, structural analysis of the PRMT-5 enzyme supports a ring-shaped PRMT-5 homodimer. The worm PRMT-5 sequence contains an insertion predicted to preclude the tetrameric architecture that underlies the heterooctomeric PRMT5-MEP50 assembly, and *C. elegans* also lack a MEP50 ortholog^17,18^.

Because *Prmt1* or *Prmt5* knockout causes early embryonic lethality in mice, organism-level analysis of PRMT-dependent arginine methylation is challenging in mammalian models^19,20^. Consistent with an essential role in vertebrate development, loss of PRMT1 in frogs results in arrested tadpole development and lethality prior to metamorphosis^21^. By contrast, *C. elegans prmt-1* and *prmt-5* single knockouts, as well as *prmt-1;prmt-5* double knockouts are viable, albeit smaller and less fertile^4^. In addition, *prmt-1* and *prmt-5* single knockout worms lack detectable ADMA and SDMA, respectively, making *C. elegans* a practical *in vivo* system for studying PRMT-dependent methylation and its physiological effects^4^. Previous studies in *C. elegans* have begun to reveal the distinct physiological roles of the individual PRMTs. For example, loss of function of *prmt-1* shortens lifespan and impairs stress tolerance by disrupting DAF-16/FoxO-mediated transcription of longevity genes^22^. By contrast, *prmt-5* deficiency does not significantly affect lifespan but instead leads to excessive germ cell apoptosis after DNA damage due to hyperactivation of p53/CEP-1 dependent cell death pathway^23^. Consistent with these findings, *prmt-5* single mutants have a lifespan close to wild type, and *prmt-1;prmt-5* double mutants exhibit similar lifespan reduction as *prmt-1* single mutants. However both *prmt-1* and *prmt-5* mutants are stress sensitive, suggesting that PRMT-5 dependent SDMA modifications are dispensable for longevity in worms even though both enzymes contribute to stress resistance^4^.

Despite these advances, the PRMT-dependent methylation landscape in *C. elegans* remains poorly defined. Most prior work has relied on organismal phenotypes and bulk measurements of methylarginine, which do not reveal which proteins and arginine sites are selectively methylated by PRMT-1 versus PRMT-5, nor how loss of each enzyme reorganizes cellular pathways at the proteome scale. Here, we address this gap by combining quantitative, site-resolved methyl-proteomics with whole-proteome profiling in wild-type worms and *prmt-1* or *prmt-5* knockout mutants. Using methyl-peptide enrichment followed by high resolution LC-MS/MS, we map enzyme dependent arginine methylation events *in vivo* and integrate these measurements with label-free proteomics to define downstream proteome remodeling. This framework enables direct comparison of *prmt-1* and *prmt-5* dependent methylation programs and identifies the biological processes most sensitive to loss of each enzyme.

## Experimental Section

### Strains

Bristol N2 (wildtype), *prmt-1* (ok2710), and *prmt-5* (gk357) null mutants obtained from the Caenorhabditis Genetics Center (CGC) were synchronized by alkaline-hypochlorite bleaching and incubated on *E. coli* OP50-seeded enriched peptone agar plates at 20 °C for 68 h. 200,000 young adults (laying eggs but eggs not hatching) were collected in triplicate, washed with ice-cold water, frozen with liquid nitrogen, and grounded in ice-cold lysis buffer (50 mM Tris-Cl, pH 7.4; 100 mM KCl; 2.5 mM MgCl₂; 0.1 % Igepal CA-630; Roche Complete protease-inhibitor cocktail) at a 1:5 animal:lysis buffer volumetric ratio. Grinding was conducted in liquid nitrogen cooled mortar and pestles.

### Lysate preparation

Proteins were precipitated in cold acetone and then resuspended in 50 mM Tris pH 7.5, 8 M urea, 1 mM activated sodium vanadate, 2.5 mM sodium pyrophosphate, 1 mM β-glycerophosphate, and 100 mM sodium phosphate. Lysates were sonicated, cleared by high-speed centrifugation, and filtered through a 0.22 µm filter. Protein concentration was measured by BCA assay, and 1 mg protein (for SCX samples) or 100 µg protein (for whole-cell lysate, WCL) was processed per sample. Next, 5 mM DTT, 25 mM iodoacetamide, and 10 mM DTT were used to reduce, alkylate, and quench protein, respectively. Lysates were diluted 4-fold in 100 mM Tris pH 8 and then trypsinized at a mass ratio of 1:100 trypsin:lysate overnight at room temperature. 5% TFA was used to quench and acidify samples to pH ∼2. For SCX enrichment, peptides were purified by using reverse phase Sep-Pak C18 cartridges (Waters) and eluted with 50% acetonitrile plus 0.1% TFA. For WCL samples, peptides were desalted using a C18 stage tip made in-house and dried by SpeedVac.

For SCX enrichment, dried 1 mg samples were resuspended in 60% acetonitrile / 40% BRUB (5 mM phosphoric acid, 5 mM boric acid, 5 mM acetic acid, pH 2.5) and incubated with high pH SCX beads (Sepax) for 30 min. After washing with washing buffer (80% acetonitrile, 20% BRUB, pH 9), four fractions with eluted with the following buffer compositions: elution buffer 1 (60% acetonitrile, 40% BRUB, pH 9), elution buffer 2 (60% acetonitrile, 40% BRUB, pH 10), and elution buffer 3 (60% acetonitrile, 40% BRUB, pH 11). Eluted fractions were dried in a SpeedVac, resuspended in 1% TFA, and desalted using an in-house stage tip with 2 mg HLB beads (Waters) and 1 layer C8 and dried by SpeedVac^24^.

### Mass Spectrometry

All liquid chromatography-mass spectrometry (LC-MS) data was collected on a nanoscale UHPLC system (EASY-nLC1200, Thermo Scientific) connected to a Q Exactive Plus hybrid quadrupole-Orbitrap mass spectrometer equipped with a nanoelectrospray source (Thermo Scientific). Peptides were separated by a reversed-phase analytical column (PepMap RSLC C18, 2 μm, 100 Å, 75 μm × 25 cm) (Thermo Scientific). For high pH SCX fractions, the flow rate was set to 300 nL/min at a gradient starting with 0% buffer B (0.1% FA, 80% acetonitrile) to 29% B in 142 min, then washed by 90% B in 10 min, and held at 90% B for 3 min. The maximum pressure was set to 1,179 bar, and the column temperature was constant at 50 °C. Dried SCX fractions were resuspended in buffer A and injected as follows: E1: 3 μL/60 μL; E2-3: 5 μL/6 μL. The effluent from the LC was directly electrosprayed into the MS. Peptides separated by the column were ionized at 2.0 kV in the positive ion mode. MS1 survey scans for data-dependent acquisition were acquired at a resolution of 70k from 350 to 1790 m/z, with a maximum injection time of 100 ms and an automatic gain control (AGC) target of 1 × 10^6^. MS/MS fragmentation of the 10 most abundant ions were analyzed at a resolution of 17.5k, AGC target of 5 × 10^4^, and a maximum injection time 240 ms, and normalized collision energies of 32 were used. Dynamic exclusion was set to 30 s and ions with charge +1 and >+6 were excluded.

### Peptide Identification

Proteome Discoverer SEQUEST (version 2.2, Thermo Scientific) was used to search MS/MS fragmentation spectra against the *in silico* tryptic digested Uniprot proteome *C. elegans* (Proteome ID UP000001940 with 26,696 sequences) and *E. coli* (strain K12i) reviewed canonical isoform downloaded May 8, 2024. For SCX samples the maximum number of missed cleavages was set to 4, and for WCL samples it was set to 2. Dynamic modifications were set to include monomethylation of arginine or lysine (R/K, +14.016 Da), dimethylation of arginine or lysine (R/K, +28.031 Da), trimethylation of lysine (K, +42.047 Da), oxidation on methionine (M, +15.995 Da), and acetylation on protein N-terminus (+42.011 Da). Fixed modification was set to carbamidomethylation on cysteine residues (C, +57.021 Da). The maximum parental mass error was set to 10 ppm, and the MS/MS mass tolerance was set to 0.02 Da. Peptides with a sequence of 7-50 amino acids were considered. For SCX samples methylation site localization was determined by the ptm-RS node in Proteome Discoverer, and only sites with localization probability greater or equal to 75% were considered. The false discovery rate (FDR) threshold was set strictly to 0.01 using the Percolator node validated by q-value.

### Methyl False Discovery Estimation

PSM and decoy PSM files were exported from Proteome Discoverer 2.2 and filtered to include only methylated and decoy PSMs. As previously described^25^, the false discovery rate (FDR) was calculated using only methyl PSMs, and target methyl PSMs were removed until a 1% FDR threshold was achieved.

### Bioinformatic Analysis

Peptide groups were exported from Proteome Discoverer 2.2 and filtered for valid values. Missing values were then imputed and normalized by in-house R code. Briefly, the data was divided into two categories under the assumption that peptides with few or no missing values exhibit missingness at random, whereas peptides that, for example, have complete data in two condition groups but three missing values in a third group exhibit missingness not at random. Missing values deemed random were imputed using KNN-TN^26^ and then combined with peptides with non-random missing values. Then peptides with non-random missing values imputed from a normal distribution down-shifted 1.8 standard deviations and with a width of 0.3 using Perseus version 2.0.6.0^27^. Imputed data was filtered for *C. elegans* peptides to exclude *E. coli* peptides, and then the peptide data was normalized by variance stabilization normalization. WCL peptide group data were summarized to the protein level by using Diffacto method^28^. Diffacto normally reports protein-level abundances at the group level (WT, *prmt1*-KO, *prmt5*-KO). We implemented a minor modification to the Diffacto codebase to also output protein-level abundances for each individual sample. PECA p-values from the Diffacto two-group comparisons (*prmt-1*-KO vs WT and *prmt-5*-KO vs WT) were used for downstream statistical analyses^29^. Q-values were calculated using the Benjamini-Hochberg method using PECA p-values. For methyl-proteomic analysis, after imputation and isoform assignment curation, only peptide groups from *C. elegans* proteome which have at least one PSMs verified by previous target-decoy FDR correction^25^ were chosen. Peptide-group abundances were normalized by VSN method. PTM annotations were derived from PSMs after target-decoy FDR filtering, then merged with the peptide-group table by sample. Methylation sites were assigned to master proteins using in-house R scripts based on peptide-level evidence.

For methyl-peptides which also had WCL protein level data, an in-house Python code based on MSstatsPTM^30^ was used to calculate p-value and t-statistics. For methyl-peptides without WCL protein level data, the Welch test was used to calculate p-values and t-statistics. After Benjamini-Hochberg FDR correction, peptides with q-value<0.05 were excluded to make a null distribution^31^ and then all peptide t-statistics were used against null distribution to get new p-value for all peptides. Then Benjamini-Hochberg was used to calculate q-values for PTM peptides.

### Neutral Loss Identification in MaxQuant

SDMA and ADMA modifications were added to MaxQuant (v2.1.3.0) as arginine dimethylation, with neutral loss masses of 31.042 (SDMA) and 45.058 (ADMA) defined in the “Neutral Loss” configuration table. RAW files were searched against Uniprot proteome *C. elegans* (Proteome ID UP000001940 with 26,696 sequences) using Trypsin/P (up to 4 missed cleavages). Carbamidomethyl (C) was set as a fixed modification, and Acetyl (Protein N-term), Oxidation (M), Methyl (KR), ADMA, and SDMA were included as variable modifications. Neutral loss annotations were extracted from msms.txt for assignment of neutral loss evidence for arginine methylation.

### Computational Assignment of Neutral Loss Evidence for Arginine Methylation

Neutral loss detection of arginine dimethylation was conducted to differentiate between ADMA and SDMA. For each MS/MS spectrum, measured parent ions and their corresponding neutral loss fragments were identified and converted to charge-adjusted neutral masses. The expected mass differences for ADMA and SDMA were compared to the observed values (31.042 for each SDMA, 45.058 for each ADMA, respectively), and acceptance was governed by a propagated error threshold derived from instrument specifications. Specifically, because both the parent ion and the neutral-loss ion contribute independently to mass measurement uncertainty (σ = 0.02 Da for each), the standard deviation of their difference is given by 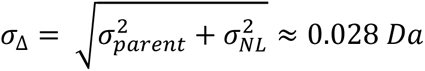. A neutral loss assignment was accepted only when the absolute residual between the observed and theoretical loss fell within this limit. For each fragment, the single best fitting theoretical loss was retained, and an additional requirement was imposed that the neutral loss peak exceeded a minimum relative intensity 0.1% threshold to ensure that only analytically robust evidence contributed to the final assignments.

Localization of dimethylation sites was performed by intersecting each validated neutral loss event with the peptide region defined by the corresponding fragment ion. When the number of candidate arginine residues within this region matched the inferred neutral loss multiplicity, the modification was assigned unambiguously to those residues. If the fragment evidence supported the presence of ADMA or SDMA but did not permit unique localization, only the aggregate ADMA/SDMA modification counts were recorded for the spectrum. The resulting annotations, comprising all accepted neutral-loss events, their localized sites, and unresolved but supported modification counts were integrated into the quantitative peptide dataset for downstream methyl-proteomic analyses.

### Motif analysis

Motif analysis was performed with pLogo^32^ by extracting 15-amino acid windows (±7 amino acids) centered on each methylated residue and uploading these sequences as the foreground. Enrichment/depletion at each flanking position was evaluated against the *C. elegans* default proteome background.

### Gene Ontology Analysis

Gene Ontology (GO) enrichment analysis was performed using ClueGO in Cytoscape. For each knockout (*prmt-1*-KO and *prmt-5*-KO), upregulated and downregulated protein sets were analyzed separately, where differential proteins were defined as those meeting |average log₂ fold change| > 0.585 (log₂1.5) and q-value < 0.05. Enrichment was tested across GO Biological Process (BP), Cellular Component (CC), and Molecular Function (MF) using GO annotations updated Sept. 20, 2025. The full set of quantified proteins (n = 2,004) was used as custom background reference. Only GO terms meeting p < 0.05 were retained and displayed. ClueGO visualization parameters included GO term fusion enabled and a GO tree interval of minimum level 3 and maximum level 8^33^.

### Data availability

All raw data have been deposited in the PRIDE database with accession number PXD074042 (reviewer access token: W25v1v3bPOwA).

## Results and Discussion

### Global analysis of protein arginine and lysine methylation in *C. elegans*

To examine how loss of PRMT-1 and PRMT-5 affects protein methylation *in vivo*, we profiled the proteome and methylome-proteome of *C. elegans* Bristol N2 (WT), *prmt-1(ok2710)*, and *prmt-5(gk357)* null mutants. Whole-worm lysates from each genotype were processed using a high-pH strong cation exchange (SCX)-LC-MS/MS workflow (Figure 1). Briefly, sonicated lysates were digested with trypsin, desalted, and split into two portions: 1 mg of peptide was subjected to high-pH SCX fractionation to enrich methyl-containing peptides^34,35^, and 250 ng was used for unfractionated whole-cell lysate (WCL) analysis. SCX fractions and WCL samples were analyzed by LC-MS/MS, and the resulting datasets were processed to generate both high-confidence methyl-site and global protein abundance profiles.

**Figure 1:**
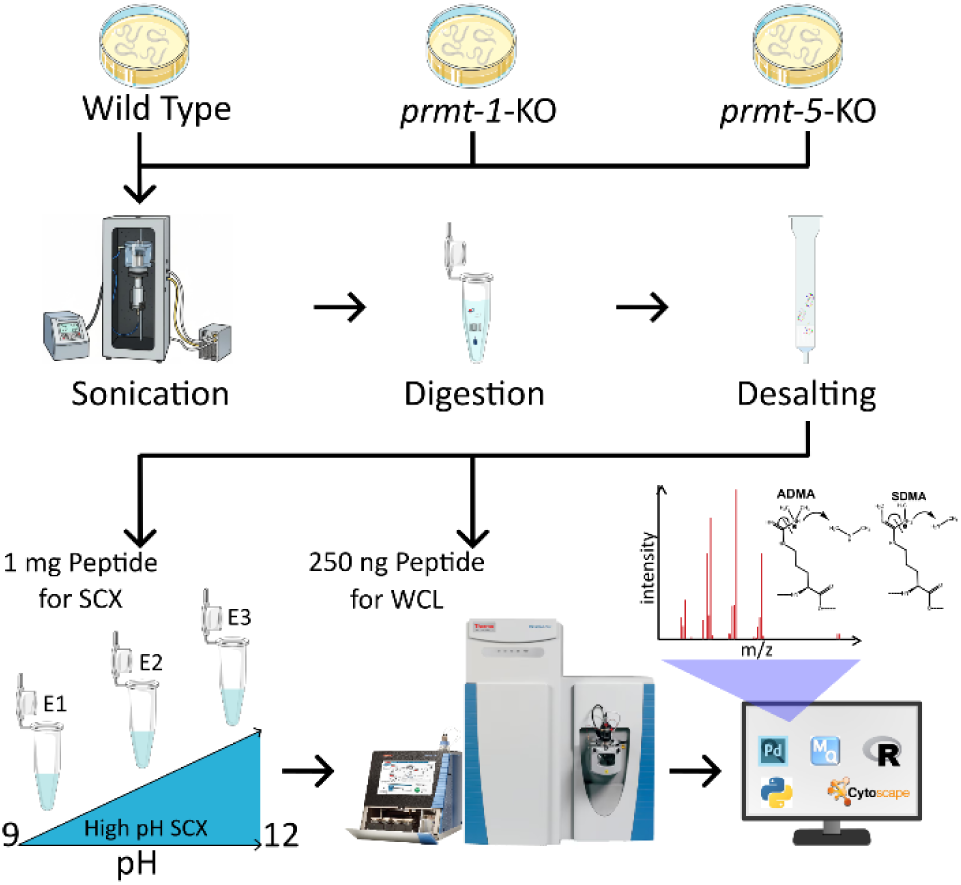
Overview of the proteomic workflow. Schematic of the workflow used to analyze the proteomic changes in wild-type, *prmt-1* knockout (*prmt-1-*KO), and *prmt-5* knockout (*prmt5*-KO) *C. elegans*. Following lysis by sonication, tryptic digestion, and desalting, samples were divided into 1) methyl-peptide analysis following enrichment of methylated peptides using high pH SCX and 2) whole cell lysate (WCL) protein abundance profiling. Samples were analyzed by LC-MS, followed by bioinformatic analysis.

Using a stringent 1% site-level methyl-FDR, we identified 347 unique methylation sites across all samples (Figure 2A, Table S1). Lysine methylation was more prevalent than arginine methylation, accounting for 74% of all methylation sites (n=257), with lysine trimethylation being the most abundant methyl-PTM identified (n=202). Among arginine methylation sites, dimethylation (n=67) was more abundant than monomethylation (n=23), consistent with the known bias of SCX enrichment^34,35^. When possible, neutral loss fragmentation was used to distinguish asymmetric (ADMA) and symmetric dimethyl-arginine (SMDA)^24^. Although not all dimethyl-arginine sites could be uniquely identified as either ADMA or SDMA, the number of confirmed ADMA sites (n=18) was much greater than confirmed SMDA sites (n=1), similar to results from mammalian cells^34,36^.

**Figure 2:**
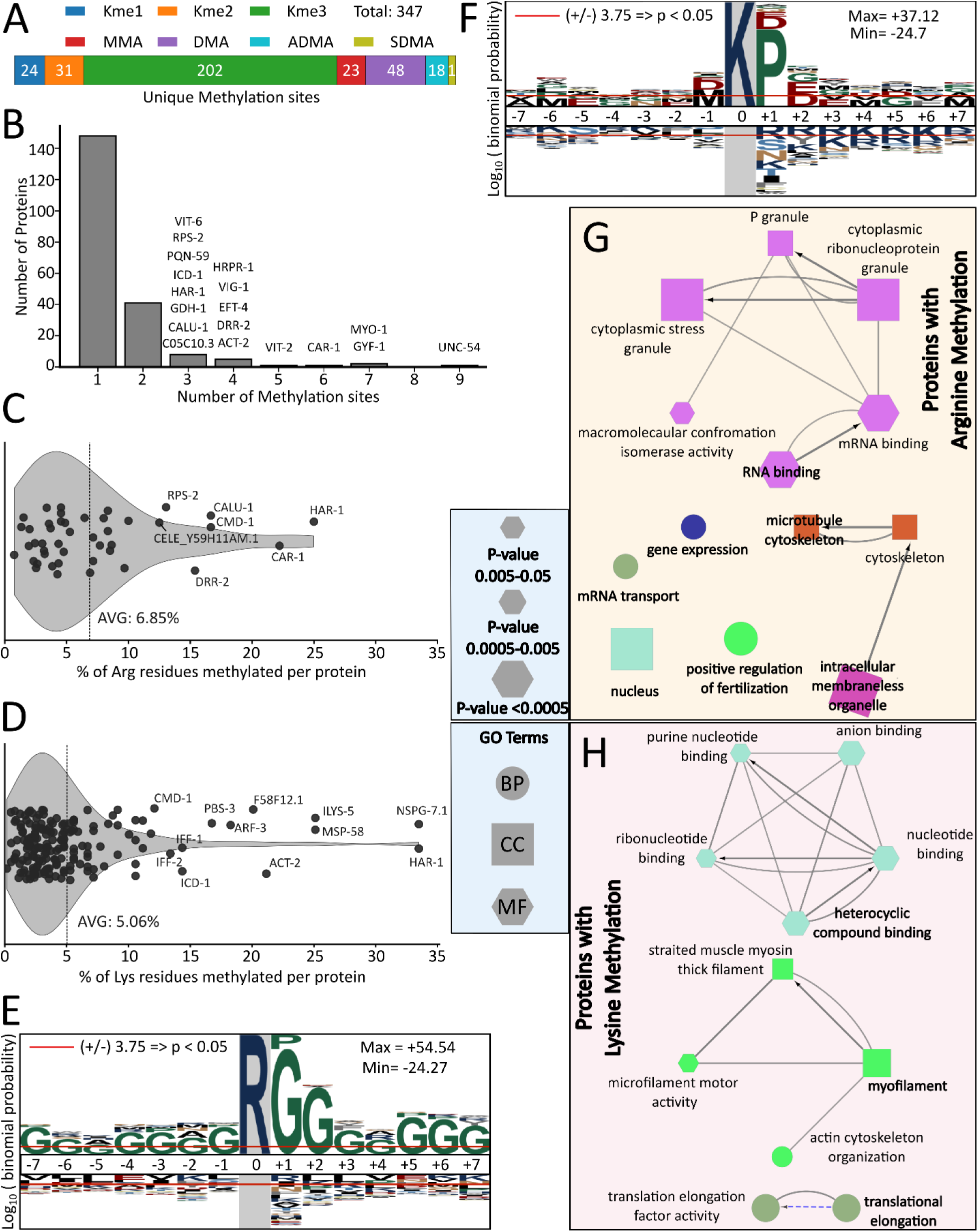
A global view of arginine and lysine methylation in *C. elegans*. A) Summary of unique methylation sites identified by LC-MS across the *C. elegans* proteome. Only high-confidence spectra passing a 1% site-level methyl-FDR were included. If a residue had multiple identified methylation forms (e.g., MMA and ADMA), each methyl proteoform was counted as a unique methylation site. ADMA and SDMA were distinguished using neutral loss fragmentation. B) Distribution of the number of methylation sites per protein for all methylated proteins. Residues with multiple identified methylation modifications were counted once. C-D) Violin plots showing the percentage of arginine (C) or lysine (D) residues that were methylated in each methylated protein. The denominator included all arginine or lysine residues present in the protein sequence, regardless of MS sequence coverage. The vertical line represents the mean methylation density across all identified proteins. E-F) Motif analysis of methylated arginine (E) and methylated lysine (F) sites identified by LC-MS. Motifs were generated using ±7-amino acid windows around methylated arginine/lysine residues compared against the entire *C. elegans* proteome. G-H) Gene ontology (GO) enrichment networks for proteins containing arginine-methylated sites (G) or lysine-methylated sites (H). In both networks, node size reflects GO-term enrichment significance (-log_10_ p-value), node color denotes functional grouping, and edges represent kappa-score connectivity indicating shared gene membership between GO terms. Biological Process (BP), Molecular Function (MF activity), and Cellular Component (CC)

Across proteins, the density of protein methylation varied substantially. While most modified proteins contained only one or two methylated residues, a subset exhibited a high density of methyl-lysine and methyl-arginine sites (Figure 2B-D). Among these, the RNA-binding protein CAR-1, UNC-54 and MYO-1, and the translational regulator GYF-1 were notable for the greatest number of methylation events (Figure 2B). CAR-1, with six methylated arginine residues out of 26 total arginine residues, ranked among the proteins with both the highest number of methylation sites and the highest percentage of methylated arginines (∼23%; Figure 2B-C). Notably, LSM14B, the human ortholog of CAR-1, has been reported to undergo extensive arginine methylation, with up to eight of 33 total arginine residues methylated (∼24%)^37^. UNC-54 exhibited the highest total number of methylation events in our dataset, with 12 distinct methylation forms distributed across nine lysine residues (Table S1). UNC-54 is orthologous to human myosin-7^38^, and seven of these nine lysine methylation sites are conserved in the human ortholog (Figure S1) although to our knowledge, none of these lysine residues in human myosin-7 have been reported to be methylated^39^.

Overall, methylated proteins contained on average 6.9% methylated arginine residues and 5.1% methylated lysine residues when normalized to the total number of arginine or lysine residues within each protein (Figure 2C-D). For comparison, a previous study of mouse muscle tissue reported that on average 5.6% of arginine residues were methylated in proteins containing methyl-arginine^40^. Of the 207 methylated proteins identified in our dataset, 163 proteins were detected exclusively with lysine methylation, while 32 proteins exhibited only arginine methylation. An additional 12 proteins contained both lysine and arginine methylation sites. Among the 12 proteins with both lysine and methylation, CMD-1 exhibits a highly conserved sequence relative to its human ortholog, calmodulin, featuring methylation at both K116 and R127 (Figure S2). Notably, both residues are conserved in the human ortholog, where K116 trimethylation and the presence of R127 have been reported to be essential for calmodulin function as a calcium-binding messenger protein^41,42^. Another highly methylated protein, MYO-1, contained six lysine methylation sites and one arginine methylation site in our dataset. Of these, five lysine sites are conserved in the human ortholog, myosin-1 (Figure S3).

Next, we used motif analysis to characterize the flanking sequence preferences for methylated arginine and lysine residues. For methyl-arginine residues, the motif was strongly enriched for RG/RGG which is the well-known glycine-arginine rich (GAR) domain (Figure 2E), consistent with established PRMT substrate preferences^43^. For methyl-lysine residues, motif analysis revealed enrichment for lysine followed by proline (KP) (Figure 2F). This KP bias may occur because C-terminal proline can prevent tryptic cleavage at lysine and because high pH SCX fractionation enriches for peptides with missed cleavages^35^.

To place these methylation sites in functional context, we next performed gene ontology (GO) enrichment on proteins containing methyl-arginine or methyl-lysine residues. Proteins bearing arginine methylation were enriched for RNA-binding functions, mRNA transport, and components of cytoplasmic ribonucleoprotein granules and stress granules, as well as nuclear and cytoskeletal processes (Figure 2G). These findings are consistent with prior studies in human cells, which have reported that arginine methylation is overrepresented among RNA-binding proteins^44^. In addition, human studies have demonstrated that arginine methylation plays a pivotal role in the formation of membraneless organelles, particularly stress granules, and in regulating their assembly and dynamics^45^. By contrast, lysine-methylated proteins were enriched for myofilament organization, motor activity, translation elongation, and nucleotide-binding functions (Figure 2H). Consistent with these results, previous studies in humans have linked lysine methylation to translation elongation^37,46^ and to myofilament organization^47^. Taken together, these data present a global picture of protein arginine and lysine methylation in *C. elegans* that is broadly consistent with the methylation profiles of mammalian cells.

### Loss of *prmt-1* alters the *C. elegans* methyl-arginine proteome

Having characterized the composition and sequence context of the protein methylome, we next quantified the effect of *prmt-1* and *prmt-5* knockout on methyl-peptide abundance (Figure 3). Only peptides with at least one peptide-spectrum match (PSM) passing a 1% methyl-peptide false discovery rate (FDR) threshold were retained, resulting in 207 quantified methylated peptides across the three SCX fractions. Duplicate peptide identifications detected across multiple fractions were collapsed by retaining the measurement with the highest observed abundance, yielding 179 unique quantified peptides for downstream analysis (Table S2). Among the 179 quantified methyl-peptides, 128 contained only methyl-lysine, 42 contained only methyl-arginine, and nine contained both methyl-arginine and methyl-lysine. Where possible, methyl-peptide abundances were normalized to changes in total protein abundance. The overall abundance of lysine methylation, as measured by the number of methylated peptide-spectrum matches (methyl-PSMs) passing the FDR threshold, was comparable across wild-type (WT), *prmt-1*-KO, and *prmt-5*-KO samples (Figure S4). In contrast, MMA was markedly reduced in *prmt-1*-KO samples, and arginine dimethylation and neutral loss confirmed ADMA were nearly undetectable in *prmt-1*-KO samples, consistent with a previous report^4^. In contrast, there were no statistically significant changes in the number of methyl-PSMs detected in *prmt-5*-KO compared to WT. These observations support that *prmt-1* knockout strongly affected protein arginine methylation but not protein lysine methylation and that *prmt-5* did not affect either protein arginine or lysine methylation.

**Figure 3.**
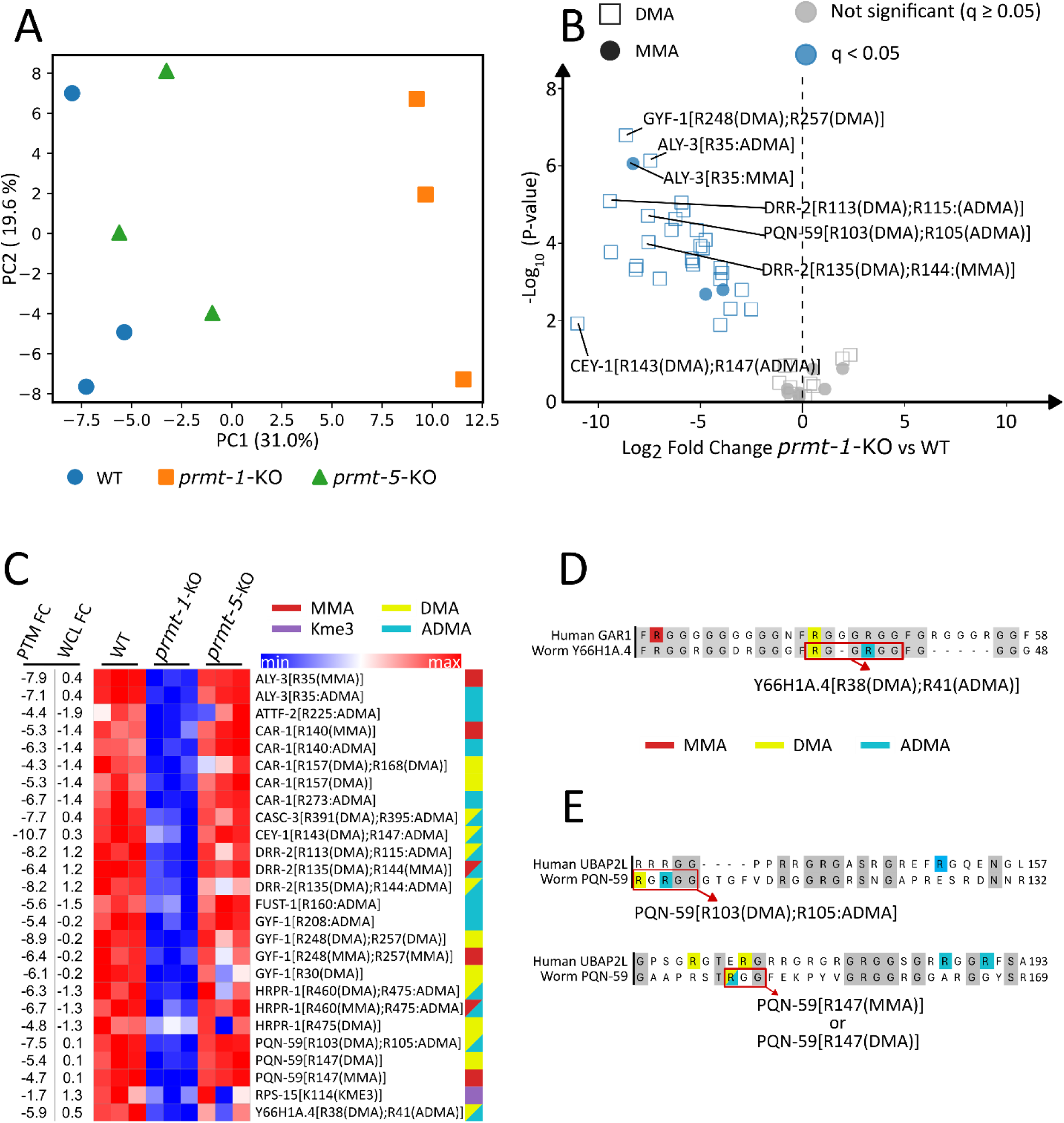
Quantitative analysis of methylated peptides in *prmt-1* knockout worms. A) Principal component analysis (PCA) of quantified methyl-peptide abundances showed separation of *prmt-1* knockout (*prmt-1*-KO) from wild-type (WT) and *prmt-5* knockout (*prmt-5*-KO) worms. B) Volcano plot comparing methyl-arginine peptide abundance between *prmt1*-KO and WT worms. Point shape denotes the identified type of methylation (MMA, DMA). Red points denote significantly upregulated peptides, and blue points denote significantly downregulated peptides (q < 0.05). C) Heatmap of the most significantly altered methylated peptides in *prmt-1*-KO worms relative to WT. The peptides shown represent those with the lowest q-values (q < 0.05), scaled by z-score across samples. At left, the log_2_ fold change (FC) values for the methyl-peptide (PTM FC) and the total protein abundance (WCL FC) are shown. The color legend at right denotes the type of methylation identified on the peptide. D-E) Sequence alignment of *C. elegans* and human orthologs with known methylation sites highlighted. Aligned residues are denoted with a gray background.

Using the 179 quantified methyl peptides, principal component analysis (PCA) revealed clear separation of *prmt-1*-KO samples from both WT and *prmt-5*-KO, while WT and *prmt-5*-KO were not distinguishable (Figure 3A). Comparing *prmt-1*-KO to wild-type (WT), 36 methyl-peptides were significantly altered at a q-value < 0.05 (Figure 3B), whereas no methyl-peptides reached this q-value threshold in *prmt-5*-KO (Figure S5). Among the 36 significantly changing methyl-peptides, 31 contained methyl-arginine, 5 contained methyl-lysine, and none contained both methyl-arginine and methyl-lysine. Notably, all 31 of the significantly changed methyl-arginine peptides were downregulated, and 28 of these significantly downregulated methyl-peptides contained DMA (90%). In addition, the fact that 31 of 42 total methyl-arginine peptides in our data set were significantly downregulated supports previous studies that PRMT1 is the dominant type I enzyme responsible for most ADMA modifications^11,12^. At several arginine residues including ALY-3 R35, CAR-1 R140, and PQN-59 R147, both the MMA and ADMA modifications were significantly decreased suggesting that PRMT-1 is responsible for successive methylation of the same residue. Among the 5 significantly changing methyl-lysine peptides, two were increased in abundance in *prmt-1-ko* samples (VIT-6[K1398(Dimethyl)], SAMS-3/4/5[K20(Trimethyl)]), but these peptides either had large decreases in total protein abundance (VIT-6) or had no total protein abundance data (SAMS-3/4/5), suggesting weak confidence in the increased methyl-peptide abundance (Figure S5). A heatmap of the quantified dataset illustrated the specificity of decreased methyl-peptide abundance in *prmt-1*-KO but not *prmt-5*-KO worms (Figure 3C). Although our results did not identify any significant changes in methyl-peptide abundance in *prmt-5*-KO worms, we cannot rule out that increased depth of methyl-peptide profiling would reveal *prmt-5* methylation targets.

Although normalized methyl-peptide data indicated a global reduction in arginine mono- and dimethylation following *prmt-1* knockout, the relationship between methylation changes and total protein abundance was not uniform across targets. For example, multiple methylation sites on GYF-1 exhibited marked decreases in *prmt-1*-KO (log_2_ fold-change ranging from -5.4 to -8.9), despite minimal changes in the corresponding master-protein abundance (log_2_ fold-change -0.2). In contrast, CAR-1 showed coordinated decreases at both the methyl-peptide (log_2_ fold-change ranging from -5.3 to -6.7) and protein levels (log_2_ fold-change -1.4), with methylation displaying a greater reduction. Conversely, proteins such as RPS-15 and DRR-2 were upregulated at the protein level in *prmt-1*-KO despite decreases in their methyl-site abundance. This suggests that methyl PTM levels and protein abundance are largely uncorrelated for these proteins.

Several proteins showing *prmt-1*-KO-specific changes in methyl-peptide abundance have been previously linked to PRMT-1 in worms and other organisms. For example, CEY-1, a Y-box-binding protein from the cold shock domain protein family, exhibited a near-complete loss of arginine dimethylation at R143 and R147 in *prmt-1*-KO worms (Figure 3C). Consistent with this observation, CEY-1 has been reported to interact with PRMT-1 in the *C. elegans* interactome^48^ and has also been shown to harbor ADMA modifications, although the specific modified residues have not previously been identified^49^. Similarly, arginine dimethylation of Y66H1A.4 (GARR-1) on R38 and R41 was significantly downregulated in *prmt-1*-KO worms. These sites are conserved in human GAR1, the ortholog of Y66H1A.4 (Figure 3D). In addition, in human cells PRMT1-mediated methylation of the GAR1 facilitates the formation and stability of H/ACA ribonucleoprotein complexes^50^, suggesting a functional role for these PTMs in worms.

Additional conserved methylation events were identified in PQN-59, the *C. elegans* ortholog of human UBAP2L, including residues R103, R105, and R147 (Figure 3E). Although PQN-59 and UBAP2L share only ∼30% overall sequence homology, they exhibit highly similar domain organization, most notably an RGG-rich region adjacent to the ubiquitin-associated (UBA) domain in both proteins^51^. In human cells, UBAP2L is a known PRMT1 substrate that regulates stress granule (SG) formation through ADMA modification^52^. While R187 and R190 have previously been proposed as ADMA sites in UBAP2L, alanine substitution of both residues did not eliminate the ADMA signal nor impair SG formation, suggesting the presence of additional functional methylation sites. Our data suggests alternative ADMA sites that may contribute to UBAP2L regulation.

For certain proteins containing repetitive RG motifs, precise alignment of methylation sites between *C. elegans* PRMT-1 targets and their human orthologs remains challenging. Nevertheless, shared interaction with PRMT1 and broadly conserved methylation patterns may provide useful clues for future functional studies. For example, HRPR-1, the *C. elegans* ortholog of human hnRNPR, displays methylation patterns that, based on sequence alignment, localize to the RGG motif of hnRNPR (Figure S6). Notably, truncating mutations within this domain of hnRNPR have recently been associated with severe developmental disorders in humans^53^. Similarly, the ADMA modification of *C. elegans* FUST-1 at R160 aligns with the region R244/R255 within the RGG1 motif of its human ortholog FUS (Figure S7). Importantly, R244 of FUS has been shown to mediate RNA binding, highlighting the potential functional relevance of this conserved methylation site^54^. Finally, CAR-1, which exhibited one of the highest numbers of methylation sites in our dataset (Figure 2B) and showed complete loss of both MMA and ADMA upon *prmt-1* knockout (Figure 3B), has a human ortholog, LSM14B (RAP55), whose localization to P-bodies is dependent on PRMT1-mediated methylation^55^. Taken together, our data reveal the widespread changes in the methyl-proteome following *prmt-1* but not *prmt-5* knockout, suggest known and novel PRMT-1 targets in *C. elegans*, and support that some methylation sites may have conserved functional significance in worms and humans.

### *prmt-1* and *prmt-5* knockout distinctly remodel the *C. elegans* proteome

To assess how *prmt-1* and *prmt-5* knockout affect broader cellular biology in *C. elegans*, we next analyzed changes in global protein abundance. Observed changes in protein abundance may, in part, reflect altered methylation of PRMT-1/PRMT-5 target proteins, but could also reflect regulatory functions of PRMT-1 and PRMT-5 that are independent of their methyltransferase activity^56,57^. Across all samples, we identified 4,018 total protein groups. After imputation and normalization, 2,004 proteins with complete quantitative values were retained for downstream analysis (Table S3-S6). Principal component analysis of these proteins revealed clear separation of *prmt-1*-KO samples with a modest separation of *prmt-5*-KO from WT samples (Figure 4A). This supports that *prmt-1* knockout more greatly affects global protein abundance profiles than does *prmt-5* knockout.

**Figure 4.**
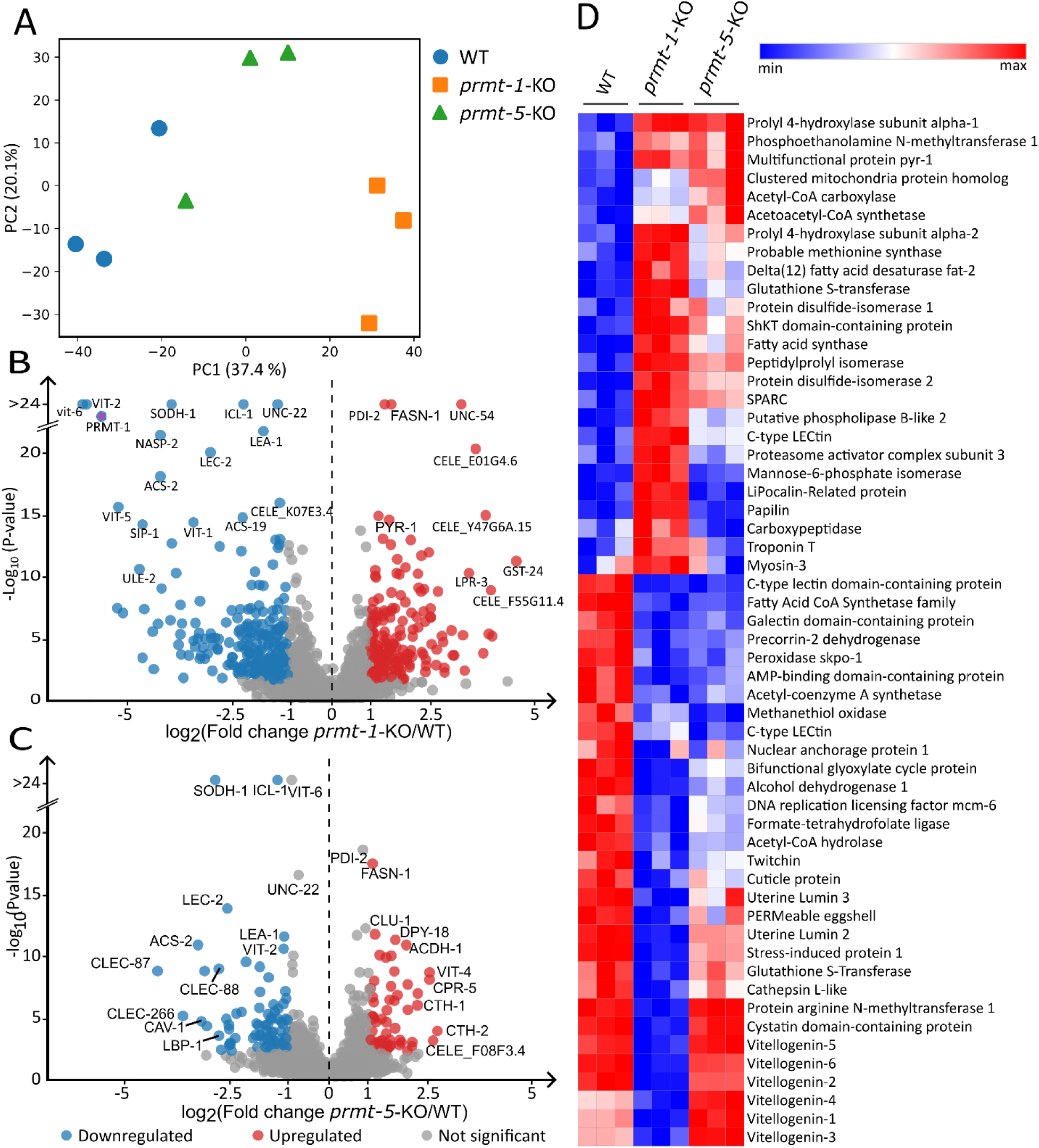
*Prmt-1* and *prmt-5* knockout remodels the *C. elegans* proteome. A) Principal component analysis (PCA) of global protein abundance profiles separated wile-type (WT), *prmt-1* knockout (*prmt-1*-KO), and *prmt-5* knockout (*prmt-5*-KO) worms. B) Volcano plot of *prmt-1*-KO versus WT protein abundances, with red and blue points indicating significantly upregulated and downregulated proteins, respectively (q < 0.05). C) Volcano plot of *prmt-5*-KO versus WT protein abundances, with red and blue points indicating significantly upregulated and downregulated proteins, respectively (q < 0.05). D) Heatmap of the most altered proteins in *prmt-1*-KO and *prmt-5*-KO worms relative to WT. Shown are the top upregulated and top downregulated proteins selected based on the lowest q-values (q < 1×10^⁻⁵^).

Indeed, pairwise comparison of *prmt-1*-KO and *prmt-5*-KO versus WT samples confirmed this finding. In *prmt-1*-KO worms, 40.5% of quantified proteins were significantly altered relative to WT (q < 0.05), and 20.8% of proteins exceeded an absolute log₂ fold-change threshold of 1 (two-fold regulation). In contrast, only 15.3% of proteins were significantly altered in the *prmt-5*-KO worms, with 6.9% surpassing the same fold-change cutoff (Figure 4B-C). Despite this difference in magnitude, *prmt-1*-KO and *prmt-5*-KO worms shared a substantial subset of significantly regulated proteins (Figure S8), and changes among shared proteins occur largely in the same direction (r = 0.56). Visualizing the proteomic abundance changes on a heatmap revealed protein groups that were upregulated in the *prmt-5*-KO but downregulated in the *prmt-1*-KO (e.g. vitellogenin 4), whereas others shift concordantly in both knockout strains (Figure 4D). The more widespread proteomic disruption observed in *prmt-1* knockout worms is consistent with prior physiological studies, which reported a significantly reduced lifespan in *prmt-1* but not *prmt-5* knockout strains^4^.

To further characterize the biological processes affected by *prmt-1* and *prmt-5* loss, we performed GO enrichment analysis (Figure 5). In *prmt-1*-KO worms, proteins reduced in abundance were predominantly associated with DNA replication, cell cycle-related pathways, and core biosynthetic or translational machinery. In contrast, proteins increased in abundance were enriched for protein folding, endoplasmic reticulum-associated functions, and amino acid and tRNA metabolic processes (Figure 5A-B). GO analysis of the *prmt-5*-KO worms displayed downregulated proteins were primarily associated with DNA replication and DNA repair pathways, whereas upregulated proteins were enriched for nucleotide- and ATP-binding functions and multiple amino acid metabolic processes (Figure 5C-D).

**Figure 5.**
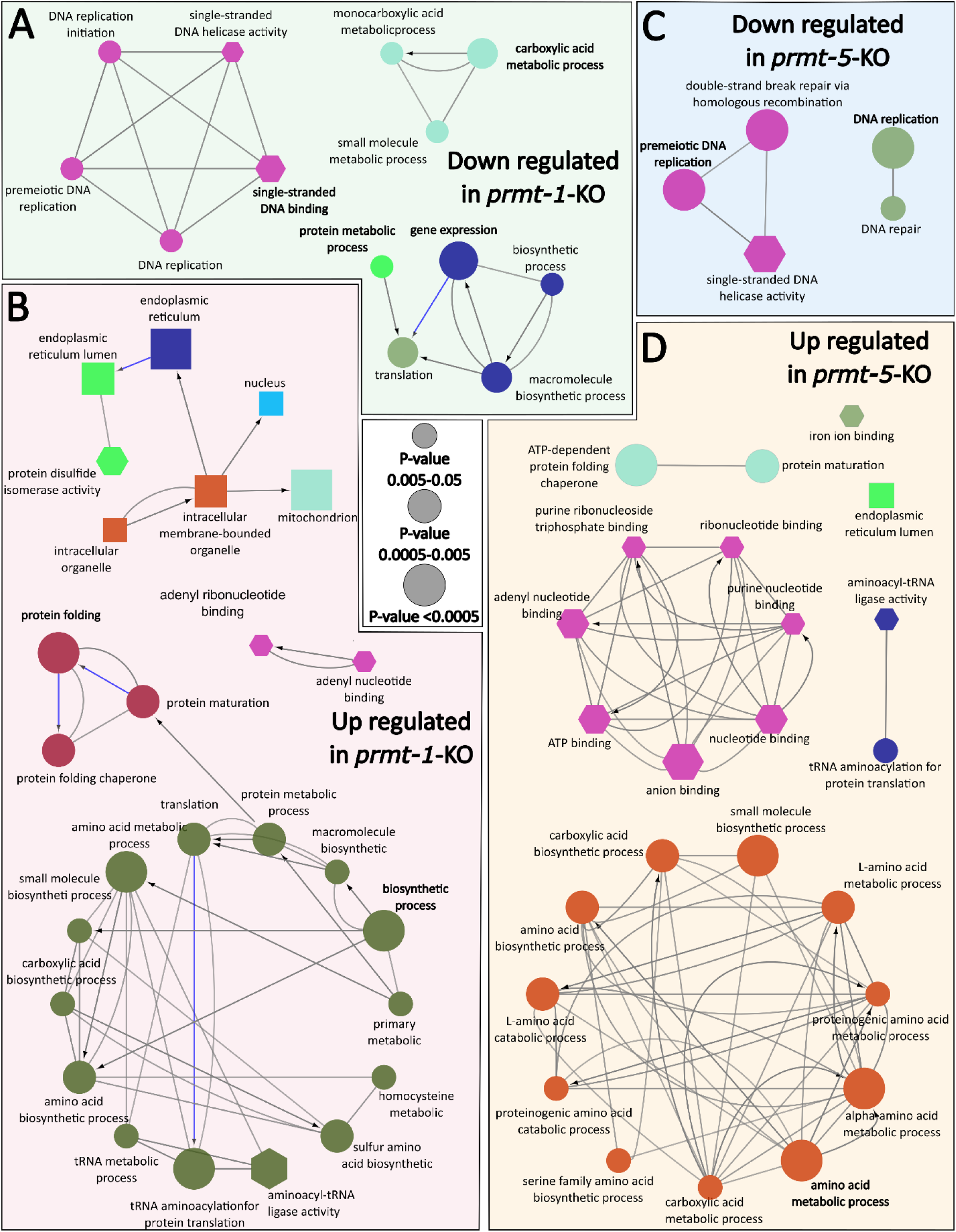
Gene ontology enrichment analysis of *prmt-1* and *prmt-5* knockout proteomes. A-B) Gene ontology (GO) enrichment analysis of the downregulated (A) and upregulated (B) proteins in *prmt-1* knockout worms (*prmt-1*-KO). Proteins included in each list met significance thresholds of |average log₂ fold change| > 0.585 (log_2_(1.5)) and q-value < 0.05 (downregulated: n = 312; upregulated: n = 402). Circles as Biological Process (BP), Hexagons as Molecular Function (MF activity), and Squares as Cellular Component (CC). All analyses were performed in ClueGO using the full set of quantified proteins (n = 2,004) as the reference background. Node size reflects the statistical significance of GO-term enrichment (-log_10_ p-value), while node color represents functional grouping. Edges represent kappa-score connectivity, indicating the degree of similarity between GO terms based on shared genes. C-D) Equivalent GO enrichment analysis for *prmt-5* knockout worms (*prmt-5*-KO) using the downregulated (n = 137) and upregulated (n = 135) protein sets.

One notable difference between *prmt-1* and *prmt-5* knockout worms was the enrichment of metabolic process-related proteins among downregulated proteins in *prmt-1* mutants. This observation is consistent with previous reports showing that *prmt-1* knockout worms exhibit a pronounced suppression of global metabolic rate and display food-avoidance behavior linked to mitochondrial dysfunction^58^. Among the most prominently affected protein groups were vitellogenins. All six vitellogenins (VIT-1 through VIT-6) were significantly downregulated in *prmt-1* knockout worms. In contrast, *prmt-5* knockout animals exhibited a mixed pattern, with VIT-1, VIT-3, VIT-4, and VIT-5 upregulated, and VIT-2 and VIT-6 significantly downregulated. Moreover, the magnitude of vitellogenin changes was substantially greater in *prmt-1* knockout worms (Figure 4B).

Vitellogenins are among the most abundant and energetically costly proteins in oviparous organisms, and their expression is tightly linked to metabolic state. One possible interpretation is that the reduced metabolic capacity of *prmt-1* knockout worms broadly suppresses vitellogenin production, whereas in *prmt-5* knockout animals, selective vitellogenin regulation may reflect a stress-adaptive response rather than global metabolic impairment. In support of this view, DAF-16^22^ and SKN-1^59^, both reported targets of PRMT-1, are known regulators of vitellogenin expression^60,61^. However, the precise mechanisms linking PRMT-1 and PRMT-5 activity to vitellogenin regulation will require further study.

### Conclusions

This study establishes an integrated, whole-animal framework for connecting PRMT-dependent methylation to downstream proteome remodeling *in vivo*. The distinct responses observed upon loss of *prmt-1* versus *prmt-5* indicate that these enzymes shape overlapping biological programs through partly different regulatory routes, consistent with contributions from both direct methylation of specific targets and broader network-level consequences. Importantly, many of the methylation sites detected occur on conserved proteins with human orthologs, providing hypotheses for conserved methylation functions and enabling future studies that translate mechanistic insights from *C. elegans* to broader PRMT biology. These findings set the stage for causal tests linking defined methylation events to protein function and phenotypic outcomes.

## Acknowledgments

This work was supported by NIH grants R21 GM144910 (NAG) and R35 GM119656 (CMP), the AACR-Bayer Innovation and Discovery Grant No. 20-80-44-GRAH (NAG), a USC Dornsife-funded Chemistry-Biology Interface fellowship (DCW), and the Viterbi School of Engineering. Some *C. elegans* strains were provided by the *Caenorhabditis* Genetics Center which is funded by NIH Office of Research Infrastructure Programs (P40 OD010440).

## Notes

### Competing Interest Statement

The authors have declared no competing interest.

