## Supplementary Figures for "LC-MS profiling of *prmt-1* and *prmt-5* knockout *C. elegans* reveals PRMT-1 substrates and global proteome remodeling"

```
sp|P02566|MYO4_CAEEL      ----MEHEKDPGWQYLRRTREQVLEDQSKPYDSKKNVWIPDPEEGYLAGEITATKGDQVT    56
sp|P12883|MYH7_HUMAN      MGDSEMAVFGAAAPYLRKSEKERLEAQTRPFDLKKDVFVPDDKQEFVKAKIVSREGGKVT    60
                                   . .  ***::.:: ** *::*:* **:*::** :: :: .:*.: :*.:**

sp|P02566|MYO4_CAEEL      IVTARGNEVTLKKELVQEMNPPKFEKTEDMSNLSFLNDASVLHNLRSRYAAMLIYTYSGL    116
sp|P12883|MYH7_HUMAN      AETEYGKTVTVKEDQVMQQNPPKFDKIEDMAMLTFLHEPAVLYNLKDRYGSWMIYTYSGL    120
                            *  *: **:*:: * : *****:* ***: *:**:: :**:**:.**.: :*******

sp|P02566|MYO4_CAEEL      FCVVINPYKRLPIYTDSCARMFMGKRKTEMPPHLFAVSDEAYRNMLQDHENQSMLITGES    176
sp|P12883|MYH7_HUMAN      FCVTVNPYKWLPVYTPEVVAAYRGKKRSEAPPHIFSISDNAYQYMLTDRENQSILITGES    180
                          ***.:**** **:** . .  : **:::* ***:*::**:**: ** *:****:******

sp|P02566|MYO4_CAEEL      GAGKTENTKKVICYFAAVGASQQEGGAEVDPNKKKVTLEDQIVQTNPVLEAFGNAKTVRN    236
sp|P12883|MYH7_HUMAN      GAGKTVNTKRVIQYFAVIAAIGDRSKKDQ--SPGKGTLEDQIIQANPALEAFGNAKTVRN    238
                          ***** ***:** ***.:.*  :..  :   .  * ******:*:**.************

sp|P02566|MYO4_CAEEL      NNSSRFGKFIRIHFNKHGRLASCDIEHYLLEKSRVIRQAPGERCYHIFYQIYSDFRPELK    296
sp|P12883|MYH7_HUMAN      DNSSRFGKFIRIHFGATGKLASADIETYLLEKSRVIFQLKAERDYHIFYQILSNKKPELL    298
                          :*************.  *:***.*** ********* *  .** ******* *: :***

sp|P02566|MYO4_CAEEL      KELLLDLPIKDYWFVAQAELIIDGIDDVEEFQLTDEAFDILNFSAVEKQDCYRLMSAHMH    356
sp|P12883|MYH7_HUMAN      DMLLITNNPYDYAFISQGETTVASIDDAEELMATDNAFDVLGFTSEEKNSMYKLTGAIMH    358
                          . **:     ** *::*.*  : .***.**:  **:***:*.*:: **:. *:* .* **

sp|P02566|MYO4_CAEEL      MGNMKFKQRPREEQAEPDGTDEAEKASNMYGIGCEEFLKALTKPRVKVGTEWVSKGQNCE    416
sp|P12883|MYH7_HUMAN      FGNMKFKLKQREEQAEPDGTEEADKSAYLMGLNSADLLKGLCHPRVKVGNEYVTKGQNVQ    418
                          :****** : **********:**:*:: : *:.. ::**.* :******.*:*:**** :

sp|P02566|MYO4_CAEEL      QVNWAVGAMAKGLYSRVFNWLVKKCNLTLDQKGIDRDYFIGVLDIAGFEIFDFNSFEQLW    476
sp|P12883|MYH7_HUMAN      QVIYATGALAKAVYERMFNWMVTRINATLETK-QPRQYFIGVLDIAGFEIFDFNSFEQLC    477
                          ** :*.**:**.:*.*:***:*.: * **: *   *:**********************

sp|P02566|MYO4_CAEEL      INFVNEKLQQFFNHHMFVLEQEEYAREGIQWVFIDFGLDLQACIELIEKPLGIISMLDEE    536
sp|P12883|MYH7_HUMAN      INFTNEKLQQFFNHHMFVLEQEEYKKEGIEWTFIDFGMDLQACIDLIEKPMGIMSILEEE    537
                          ***.******************** :***:*.*****:******:*****:**:*:*:**

sp|P02566|MYO4_CAEEL      CIVPKATDLTLASKLVDQHLGKHPNFEKPKPPKGKQGEAHFAMRHYAGTVRYNCLNWLEK    596
sp|P12883|MYH7_HUMAN      CMFPKATDMTFKAKLFDNHLGKSANFQKPRNIKGK-PEAHFSLIHYAGIVDYNIIGWLQK    596
                          *:.*****:*: :**.*:****  **:**:  ***  ****:: **** * ** :.**:*

sp|P02566|MYO4_CAEEL      NKDPLNDTVVSAMKQSKGNDLLVEIWQDYTTQEEAAAKAKEGGGGGKKKGKSGSFMTVSM    656
sp|P12883|MYH7_HUMAN      NKDPLNETVVGLYQKS-SLKLLSTLFANYAGADAPIE----KGKG--KAKKGSSFQTVSA    649
                          ******:***.  ::* . .**  :: :*:  :         * *  *  *..** ***

sp|P02566|MYO4_CAEEL      LYRESLNNLMTMLNKTHPHFIRCIIPNEKKQSGMIDAALVLNQLTCNGVLEGIRICRKGF    716
sp|P12883|MYH7_HUMAN      LHRENLNKLMTNLRSTHPHFVRCIIPNETKSPGVMDNPLVMHQLRCNGVLEGIRICRKGF    709
                          *:**.**:*** *..*****:*******.*. *::*  **::** ***************

sp|P02566|MYO4_CAEEL      PNRTLHPDFVQRYAILAAKEAKSDD--DKKKCAEAIMSKLVNDGSLSEEMFRIGLTKVFF    774
sp|P12883|MYH7_HUMAN      PNRILYGDFRQRYRILNPAAIPEGQFIDSRKGAE----KLLSSLDIDHNQYKFGHTKVFF    765
                          *** *: ** *** **      ..:  *.:* **    **:.. .:..: :::* *****

sp|P02566|MYO4_CAEEL      KAGVLAHLEDIRDEKLATILTGFQSQIRWHLGLKDRKRRMEQRAGLLIVQRNVRSWCTLR    834
sp|P12883|MYH7_HUMAN      KAGLLGLLEEMRDERLSRIITRIQAQSRGVLARMEYKKLLERRDSLLVIQWNIRAFMGVK    825
                          ***:*. **::***:*: *:* :*:* *  *.  : *: :*:* .**::* *:*::  ::

sp|P02566|MYO4_CAEEL      TWEWFKLYGKVKPMLKAGKEAEELEKINDKVKALEDSLAKEEKLRKELEESSAKLVEEKT    894
sp|P12883|MYH7_HUMAN      NWPWMKLYFKIKPLLKSAEREKEMASMKEEFTRLKEALEKSEARRKELEEKMVSLLQEKN    885
                          .* *:*** *:**:**:.:. :*: .::::.. *:::* *.*  ******. ..*::**.

sp|P02566|MYO4_CAEEL      SLFTNLESTKTQLSDAEERLAKLEAQQKDASKQLSELNDQLADNEDRTADVQRAKKKIEA    954
```

```
sp|P12883|MYH7_HUMAN      DLQLQVQAEQDNLADAEERCDQLIKNKIQLEAKVKEMNERLEDEEEMNAELTAKKRKLED    945
                          .*  :::: : :*:*****  :*  :: : . ::.*:*::* *:*: .*::   *:*:*

sp|P02566|MYO4_CAEEL      EVEALKKQIQDLEMSLRKAESEKQSKDHQIRSLQDEMQQQDEAIAKLNKEKKHQEEINRK    1014
sp|P12883|MYH7_HUMAN      ECSELKRDIDDLELTLAKVEKEKHATENKVKNLTEEMAGLDEIIAKLTKEKKALQEAHQQ    1005
                          * . **::*:***::* *.*.**::.:::::.* :**   ** ****.****  :* :::

sp|P02566|MYO4_CAEEL      LMEDLQSEEDKGNHQNKVKAKLEQTLDDLEDSLEREKRARADLDKQKRKVEGELKIAQEN    1074
sp|P12883|MYH7_HUMAN      ALDDLQAEEDKVNTLTKAKVKLEQQVDDLEGSLEQEKKVRMDLERAKRKLEGDLKLTQES    1065
                           ::***:**** *  .*.*.**** :****.***:**:.* **:: ***:**:**::**.

sp|P02566|MYO4_CAEEL      IDESGRQRHDLENNLKKKESELHSVSSRLEDEQALVSKLQRQIKDGQSRISELEEELENE    1134
sp|P12883|MYH7_HUMAN      IMDLENDKQQLDERLKKKDFELNALNARIEDEQALGSQLQKKLKELQARIEELEEELEAE    1125
                          * :  .::::*::.****: **:::.:*:****** *:**:::*: *:**.******* *

sp|P02566|MYO4_CAEEL      RQSRSKADRAKSDLQRELEELGEKLDEQGGATAAQVEVNKKREAELAKLRRDLEEANMNH    1194
sp|P12883|MYH7_HUMAN      RTARAKVEKLRSDLSRELEEISERLEEAGGATSVQIEMNKKREAEFQKMRRDLEEATLQH    1185
                          * :*:*.:: :***.*****:.*:*:* ****:.*:*:*******: *:*******.::*

sp|P02566|MYO4_CAEEL      ENQLGGLRKKHTDAVAELTDQLDQLNKAKAKVEKDKAQAVRDAEDLAAQLDQETSGKLNN    1254
sp|P12883|MYH7_HUMAN      EATAAALRKKHADSVAELGEQIDNLQRVKQKLEKEKSEFKLELDDVTSNMEQIIKAKANL    1245
                          *   ..*****:*:**** :*:*:*::.* *:**:*::   : :*::::::*  ..* *

sp|P02566|MYO4_CAEEL      EKLAKQFELQLTELQSKADEQSRQLQDFTSLKGRLHSENGDLVRQLEDAESQVNQLTRLK    1314
sp|P12883|MYH7_HUMAN      EKMCRTLEDQMNEHRSKAEETQRSVNDLTSQRAKLQTENGELSRQLDEKEALISQLTRGK    1305
                          **:.: :* *:.* :***:* .*.::*:** :.:*::***:* ***:: *: :.**** *

sp|P02566|MYO4_CAEEL      SQLTSQLEEARRTADEEARERQTVAAQAKNYQHEAEQLQESLEEEIEGKNEILRQLSKAN    1374
sp|P12883|MYH7_HUMAN      LTYTQQLEDLKRQLEEEVKAKNALAHALQSARHDCDLLREQYEEETEAKAELQRVLSKAN    1365
                             *.***: :*  :**.: ::::*   :. :*:.: *:*. *** *.* *: * *****

sp|P02566|MYO4_CAEEL      ADIQQWKARFEGEGLLKADELEDAKRRQAQKINELQEALDAANSKNASLEKTKSRLVGDL    1434
sp|P12883|MYH7_HUMAN      SEVAQWRTKYETDAIQRTEELEEAKKKLAQRLQEAEEAVEAVNAKCSSLEKTKHRLQNEI    1425
                          ::: **::::* :.: :::***:**:: **:::* :**::*.*:* :****** ** .::

sp|P02566|MYO4_CAEEL      DDAQVDVERANGVASALEKKQKGFDKIIDEWRKKTDDLAAELDGAQRDLRNTSTDLFKAK    1494
sp|P12883|MYH7_HUMAN      EDLMVDVERSNAAAAALDKKQRNFDKILAEWKQKYEESQSELESSQKEARSLSTELFKLK    1485
                          :*  *****:*..*:**:***:.****: **::* ::  :**:.:*:: *. **:*** *

sp|P02566|MYO4_CAEEL      NAQEELAEVVEGLRRENKSLSQEIKDLTDQLGEGGRSVHEMQKIIRRLEIEKEELQHALD    1554
sp|P12883|MYH7_HUMAN      NAYEESLEHLETFKRENKNLQEEISDLTEQLGSSGKTIHELEKVRKQLEAEKMELQSALE    1545
                          ** **  * :* ::****.*.:**.***:***..*:::**::*: ::** ** *** **:

sp|P02566|MYO4_CAEEL      EAEAALEAEESKVLRAQVEVSQIRSEIEKRIQEKEEEFENTRKNHARALESMQASLETEA    1614
sp|P12883|MYH7_HUMAN      EAEASLEHEEGKILRAQLEFNQIKAEIERKLAEKDEEMEQAKRNHLRVVDSLQTSLDAET    1605
                          ****:** **.*:****:*..**::***::: **:**:*::::** *.::*:*:**::*:

sp|P02566|MYO4_CAEEL      KGKAELLRIKKKLEGDINELEIALDHANKANADAQKNLKRYQEQVRELQLQVEEEQRNGA    1674
sp|P12883|MYH7_HUMAN      RSRNEALRVKKKMEGDLNEMEIQLSHANRMAAEAQKQVKSLQSLLKDTQIQLDDAVRAND    1665
                          :.: * **:***:***:**:** *.***:  *:***::*  *. ::: *:*:::  * .

sp|P02566|MYO4_CAEEL      DTREQFFNAEKRATLLQSEKEELLVANEAAERARKQAEYEAADARDQANEANAQVSSLTS    1734
sp|P12883|MYH7_HUMAN      DLKENIAIVERRNNLLQAELEELRAVVEQTERSRKLAEQELIETSERVQLLHSQNTSLIN    1725
                          * :*::  .*:* .***:* *** .. * :**:** ** *  :: ::.:  ::* :** .

sp|P02566|MYO4_CAEEL      AKRKLEGEIQAIHADLDETLNEYKAAEERSKKAIADATRLAEELRQEQEHSQHVDRLRKG    1794
sp|P12883|MYH7_HUMAN      QKKKMDADLSQLQTEVEEAVQECRNAEEKAKKAITDAAMMAEELKKEQDTSAHLERMKKN    1785
                           *:*::.::. ::::::*:::* : ***::****:**: :****::**: * *::*::*.

sp|P02566|MYO4_CAEEL      LEQQLKEIQVRLDEAEAAALKGGKKVIAKLEQRVRELESELDGEQRRFQDANKNLGRADR    1854
sp|P12883|MYH7_HUMAN      MEQTIKDLQHRLDEAEQIALKGGKKQLQKLEARVRELENELEAEQKRNAESVKGMRKSER    1845
```

```
                              :** :*::* ******  ******* : *** ******.**:.**:*  :: *.: :::*

sp|P02566|MYO4_CAEEL      RVRELQFQVDEDKKNFERLQDLIDKLQQKLKTQKKQVEEAEELANLNLQKYKQLTHQLED    1914
sp|P12883|MYH7_HUMAN      RIKELTYQTEEDRKNLLRLQDLVDKLQLKVKAYKRQAEEAEEQANTNLSKFRKVQHELDE    1905
                          *::** :*.:**:**: *****:**** *:*: *:*.***** ** **.*:::: *:*::

sp|P02566|MYO4_CAEEL      AEERADQAENSLSKMRSKSRASASVAPGLQSSASAAVIRSPSRARASDF       1963
sp|P12883|MYH7_HUMAN      AEERADIAESQVNKLRAKSRDIGTKGL--NEE-----------------       1935
                          ****** **..:.*:*:***  .: .   :..
```

**Figure S1: Conservation of UNC-54 methylation sites in human MYH7.**

Protein sequences of *C. elegans* UNC-54 (MYO-4_CAEEL) and human myosin-7 (MYH7_HUMAN) were aligned using Clustal Omega v1.2.4. Methylation sites detected on UNC-54 were mapped onto the alignment to identify corresponding residues in MYH7. The following UNC-54 lysine sites align to MYH7 positions: K185→K189, K381→K383, K571→K572, K644→K637, K647→K640, K686→K679, and K1626→K1617. Two sites did not map to a corresponding lysine in MYH7 (K51 and K645)

```
sp|P0DP25|CALM3_HUMAN      MADQLTEEQIAEFKEAFSLFDKDGDGTITTKELGTVMRSLGQNPTEAELQDMINEVDADG   60
sp|P0DP24|CALM2_HUMAN      MADQLTEEQIAEFKEAFSLFDKDGDGTITTKELGTVMRSLGQNPTEAELQDMINEVDADG   60
sp|P0DP23|CALM1_HUMAN      MADQLTEEQIAEFKEAFSLFDKDGDGTITTKELGTVMRSLGQNPTEAELQDMINEVDADG   60
sp|O16305|CALM_CAEEL       MADQLTEEQIAEFKEAFSLFDKDGDGTITTKELGTVMRSLGQNPTEAELQDMINEVDADG   60
                           ************************************************************

sp|P0DP25|CALM3_HUMAN      NGTIDFPEFLTMMARKMKDTDSEEEIREAFRVFDKDGNGYISAAELRHVMTNLGEKLTDE   120
sp|P0DP24|CALM2_HUMAN      NGTIDFPEFLTMMARKMKDTDSEEEIREAFRVFDKDGNGYISAAELRHVMTNLGEKLTDE   120
sp|P0DP23|CALM1_HUMAN      NGTIDFPEFLTMMARKMKDTDSEEEIREAFRVFDKDGNGYISAAELRHVMTNLGEKLTDE   120
sp|O16305|CALM_CAEEL       NGTIDFPEFLTMMARKMKDTDSEEEIREAFRVFDKDGNGFISAAELRHVMTNLGEKLTDE   120
                           ***************************************:********************

sp|P0DP25|CALM3_HUMAN      EVDEMIREADIDGDGQVNYEEFVQMMTAK    149
sp|P0DP24|CALM2_HUMAN      EVDEMIREADIDGDGQVNYEEFVQMMTAK    149
sp|P0DP23|CALM1_HUMAN      EVDEMIREADIDGDGQVNYEEFVQMMTAK    149
sp|O16305|CALM_CAEEL       EVDEMIREADIDGDGQVNYEEFVTMMTTK    149
                           *********************** ***:*
```

**Figure S2: Conservation of calmodulin methylation sites in C. elegans CMD-1 and human calmodulin.**

Multiple sequence alignment of C. elegans CMD-1 (CALM_CAEEL) with human calmodulin isoforms (CALM1, CALM2, and CALM3) was performed using Clustal Omega v1.2.4. The alignment highlights the strong sequence conservation of calmodulin and was used to map methylation sites detected in CMD-1. CMD-1 methylation at K116 and R127 is shown, and both residues are conserved in human orthologs. Consistent with prior reports, K116 trimethylation and the presence of R127 have been implicated as important for calmodulin function as a calcium-binding messenger protein.

```
sp|P12882|MYH1_HUMAN      MSSDSEMAIFG---EAAPFLRKSERERIEAQNKPFDAKTSVFVVDPKESFVKATVQSREG    57
sp|P02567|MYO1_CAEEL      ------MSLEHEKDPGWQYLKRSREQQLADQSRPYDSKKNVWIPDAEEGYIEGVIKGPGP    54
                                *::      .  :*::*..:::  *.:*:*:*..*:: * :*.:::..::.

sp|P12882|MYH1_HUMAN      G-KVTAKTEAGATVTVKDDQVFPMNPPKYDKIEDMAMMTHLHEPAVLYNLKERYAAWMIY    116
sp|P02567|MYO1_CAEEL      KADTVIVTAGGKDVTLKKDIVQEVNPPKFEKTEDMSNLTFLNDASVLWNLRSRYAAMLIY    114
                            ...   * .*  **:*.* *  :****::* ***: :*.*:: :**:**:.**** :**

sp|P12882|MYH1_HUMAN      TYSGLFCVTVNPYKWLPVYNAEVVTAYRGKKRQEAPPHIFSISDNAYQFMLTDRENQSIL    176
sp|P02567|MYO1_CAEEL      TYSGLFCVVINPYKRLPIYTDSVARMFMGKRRTEMPPHLFAVSDQAYRYMLQDHENQSML    174
                          ********.:**** **:*. .*.  : **:* * ***:*::**:**::** *:****:*

sp|P12882|MYH1_HUMAN      ITGESGAGKTVNTKRVIQYFATIAVTGEKKKEEVTSGKMQGTLEDQIISANPLLEAFGNA    236
sp|P02567|MYO1_CAEEL      ITGESGAGKTENTKKVICYFATVGAS---QKAALKEGEKEVTLEDQIVQTNPVLEAFGNA    231
                          ********** ***:** ****:..:   :*  :..*: : ******:.:**:*******

sp|P12882|MYH1_HUMAN      KTVRNDNSSRFGKFIRIHFGTTGKLASADIETYLLEKSRVTFQLKAERSYHIFYQIMSNK    296
sp|P02567|MYO1_CAEEL      KTVRNNNSSRFGKFIRIHFNKHGTLASCDIEHYLLEKSRVIRQAPGERCYHIFYQIYSDF    291
                          *****:*************.. *.***.*** ********  *  .**.******* *:

sp|P12882|MYH1_HUMAN      KPDLIEMLLITTNPYDYAFVSQGEITVPSIDDQEELMATDSAIEILGFTSDERVSIYKLT    356
sp|P02567|MYO1_CAEEL      KPQLRDELLLNHPISNYWFVAQAELLIDGIDDTEEFQLTDEAFDVLKFSPTEKMDCYRLM    351
                          **:* : **:.    :* **:*.*: : .*** **:  **.*:::* *:  *::. *:*

sp|P12882|MYH1_HUMAN      GAVMHYGNMKFKQKQREEQAEPDGTEVADKAAYLQNLNSADLLKALCYPRVKVGNEYVTK    416
sp|P02567|MYO1_CAEEL      SAHMHMGNMKFKQRPREEQAEPDGQDEAERACNMYGIDVDQFLKALVSPRVKVGTEWVSK    411
                          .* ** *******: ********* : *::*. : .::  ::****  ******.*:*:*

sp|P12882|MYH1_HUMAN      GQTVQQVYNAVGALAKAVYDKMFLWMVTRINQQLDTK-QPRQYFIGVLDIAGFEIFDFNS    475
sp|P02567|MYO1_CAEEL      GQNVDQVHWAIGAMAKGLYARVFHWLVKKCNLTLDQKGIDRDYFIGVLDIAGFEIFDFNS    471
                          **.*:**: *:**:**.:* ::* *:*.: *  ** *   *:*****************

sp|P12882|MYH1_HUMAN      LEQLCINFTNEKLQQFFNHHMFVLEQEEYKKEGIEWTFIDFGMDLAACIELIEKPMGIFS    535
sp|P02567|MYO1_CAEEL      FEQLWINFVNEKLQQFFNHHMFVLEQEEYAREGIQWTFIDFGLDLQACIELIEKPLGIIS    531
                          :*** ***.******************** :***:*******:** *********:**:*

sp|P12882|MYH1_HUMAN      ILEEECMFPKATDTSFKNKLYEQHLGKSNNFQKPKPAKGKP-EAHFSLIHYAGTVDYNIA    594
sp|P02567|MYO1_CAEEL      MLDEECIVPKATDMTLAQKLTDQHLGKHPNFEKPKPPKGKQGEAHFAMRHYAGTVRYNVL    591
                          :*:***:.***** :: :** :*****  **:**** ***  ****:: ****** **:

sp|P12882|MYH1_HUMAN      GWLDKNKDPLNETVVGLYQKSAMKTLALLFV------GATGAEAEAGGGKKGGKKKGSSF    648
sp|P02567|MYO1_CAEEL      NWLEKNKDPLNDTVVSVMKASKKNDLLVEIWQDYTTQEEAAAAAKAGGGRKGGK--SGSF    649
                          .**:*******:***.: : *  : * : :         :.* *:****:****  ..**

sp|P12882|MYH1_HUMAN      QTVSALFRENLNKLMTNLRSTHPHFVRCIIPNETKTPGAMEHELVLHQLRCNGVLEGIRI    708
sp|P02567|MYO1_CAEEL      MTVSMMYRESLNKLMTMLHKTHPHFIRCIIPNEKKQSGMIDAALVLNQLTCNGVLEGIRI    709
                           *** ::**.****** *:.*****:*******.*  * ::  ***:** **********

sp|P12882|MYH1_HUMAN      CRKGFPSRILYADFKQRYKVLNASAIPEGQFIDS--KKASEKLLGSIDIDHTQYKFGHTK    766
sp|P02567|MYO1_CAEEL      CRKGFPNRTQHPDFVQRYAILAAKEAKSSDDMKTCAGAILQALINQKQLNDEQFRIGHTK    769
                          ******.*  : ** *** :* *.   ..: :.:      : *:.. :::. *:::****

sp|P12882|MYH1_HUMAN      VFFKAGLLGLLEEMRDEKLAQLITRTQAMCRGFLARVEYQKMVERRESIFCIQYNVRAFM    826
sp|P02567|MYO1_CAEEL      VFFKAGVVAHIEDLRDDKLNQIITGFQSAIRWYTATADAGARRKQLNSYIILQRNIRSWC    829
                          ******::. :*::**:** *:**  *:  * : * .:     :: :* : :* *:*::

sp|P12882|MYH1_HUMAN      NVKHWPWMKLYFKIKPLLKSAETEKEMANMKEEFEKTKEELAKTEAKRKELEEKMVTLMQ    886
sp|P02567|MYO1_CAEEL      VLRTWDWFLLFGKLRPQLKCGKMAEEMIKMAEEQKVLEAEAKKAESARKSQEEAYAKLSA    889
                           :: * *: *: *::* **..:  :** :* ** :  : *  *:*: **. **  ..*

sp|P12882|MYH1_HUMAN      EKNDLQLQVQAEADSLADAEERCDQLIKTKIQLEAKIKEVTERAEDEEEINAELTAKKRK    946
```

```
sp|P02567|MYO1_CAEEL      ERSKLLEALELTQGGSAAIEEKLTRLNSARQEVEKSLNDANDRLSEHEEKNADLEKQRRK    949
                          *:..*   ::   .. *  **:  :* .:: ::* .:::..:* .:.** **:*  ::**

sp|P12882|MYH1_HUMAN      LEDECSELKKDIDDLELTLAKVEKEKHATENKVKNLTEEMAGLDETIAKLTKEKKALQEA    1006
sp|P02567|MYO1_CAEEL      AQQEVENLKKSIEAVDGNLAKSLEEKAAKENQIHSLQDEMNSQDETIGKINKEKKLLEEN    1009
                           ::* .:***.*: :: .***  :** *.**:::.* :** . ****.*:.**** *:*

sp|P12882|MYH1_HUMAN      HQQTLDDLQAEEDKVNTLTKAKIKLEQQVDDLEGSLEQEKKIRMDLERAKRKLEGDLKLA    1066
sp|P02567|MYO1_CAEEL      NRQLVDDLQAEEAKQAQANRLRGKLEQTLDEMEEAVEREKRIRAETEKSKRKVEGELKGA    1069
                          ::* :******* *    .: : **** :*::* ::*:**:** : *::***:**:** *

sp|P12882|MYH1_HUMAN      QESTMDIENDKQQLDEKLKKKEFEMSGLQSKIEDEQALGMQLQKKIKELQARIEELEEEI    1126
sp|P02567|MYO1_CAEEL      QETIDELSAIKLETDASLKKKEADIHALGVRIEDEQALANRLTRQSKENAQRIIEIEDEL    1129
                          **:  ::.  * : * .***** :: .*  :*******. :* :: **   ** *:*:*:

sp|P12882|MYH1_HUMAN      EAERASRAKAEKQRSDLSRELEEISERLEEAGGATSAQIEMNKKREAEFQKMRRDLEEAT    1186
sp|P02567|MYO1_CAEEL      EHERQSRSKADRARAELQRELDELNERLDEQNKQLEIQQDNNKKKDSEIIKFRRDLDEKN    1189
                          * ** **:**:: *::*.***:*:.***:* .    . * : ***:::*: *:****:* .

sp|P12882|MYH1_HUMAN      LQHEATAATLRKKHADSVAELGEQIDNLQRVKQKLEKEKSEMKMEIDDLASNMETVSKAK    1246
sp|P02567|MYO1_CAEEL      MANEDQMAMIRRKNNDQISALTNTLDALQKSKAKIEKEKGVLQKELDDINAQVDQETKSR    1249
                          : :*   * :*:*: *.:: * : :* **: * *:****. :: *:**: ::::  :*::

sp|P12882|MYH1_HUMAN      GNLEKMCRALEDQLSEIKTKEEEQQRLINDLTAQRARLQTESGEYSRQLDEKDTLVSQLS    1306
sp|P02567|MYO1_CAEEL      VEQERLAKQYEIQVAELQQKVDEQSRQIGEYTSTKGRLSNDNSDLARQVEELEIHLATIN    1309
                           : *::.:  * *::*:: * :**.* *.: *: :.**..:..: :**::* :  :: :.

sp|P12882|MYH1_HUMAN      RGKQAFTQQIEELKRQLEEEIKAKSALAHALQSSRHDCDLLREQYEEEQEAKAELQRAMS    1366
sp|P02567|MYO1_CAEEL      RAKTAFSSQLVEAKKAAEDELHERQEFHAACKNLEHELDQCHELLEEQINGKDDIQRQLS    1369
                          *.* **:.*: * *:  *:*:: :. :  * :. .*: *  :*  **: :.* ::** :*

sp|P12882|MYH1_HUMAN      KANSEVAQWRTKYETDAIQRTEELEEAKKKLAQRLQDAEEHVEAVNAKCASLEKTKQRLQ    1426
sp|P02567|MYO1_CAEEL      RINSEISQWKARYEGEGLVGSEELEELKRKQMNRVMDLQEALSAAQNKVISLEKAKGKLL    1429
                          : ***::**:::** :.:  :***** *:*  :*: * :* :.*.: *  ****:* :*

sp|P12882|MYH1_HUMAN      NEVEDLMIDVERTNAACAALDKKQRNFDKILAEWKQKCEETHAELEASQKESRSLSTELF    1486
sp|P02567|MYO1_CAEEL      AETEDARSDVDRHLTVIASLEKKQRAFDKIVDDWKRKVDDIQKEIDATTRDSRNTSTEVF    1489
                           *.**   **:*  :. *:*:**** ****: :**:* :: : *::*: ::**. ***:*

sp|P12882|MYH1_HUMAN      KIKNAYEESLDQLETLKRENKNLQQEISDLTEQIAEGGKRIHELEKIKKQVEQEKSELQA    1546
sp|P02567|MYO1_CAEEL      KLRSSMDNLSEQIETLRRENKIFSQEIRDINEQITQGGRTYQEVHKSVRRLEQEKDELQH    1549
                          *::.: ::  :*:***:**** :.*** *:.***::**:  :*:.*  :::****.***

sp|P12882|MYH1_HUMAN      ALEEAEASLEHEEGKILRIQLELNQVKSEVDRKIAEKDEEIDQMKRNHIRIVESMQSTLD    1606
sp|P02567|MYO1_CAEEL      ALDEAEAALEAEESKVLRLQIEVQQIRSEIEKRIQEKEEEFENTRKNHQRALESIQASLE    1609
                          **:****:** **.*:**:*:*::*::**::**::::* **:**::: ::** * :**:*::*:

sp|P12882|MYH1_HUMAN      AEIRSRNDAIRLKKKMEGDLNEMEIQLNHANRMAAEALRNYRNTQAILKDTQLHLDDALR    1666
sp|P02567|MYO1_CAEEL      TEAKSKAELARAKKKLETDINQLEIALDHANKANVDAQKNLKKLFDQVKELQGQVDDEQR    1669
                          :* :*: :  * ***:* *:*::** *:***:  .:* :* ::    :*: * ::**  *

sp|P12882|MYH1_HUMAN      SQEDLKEQLAMVERRANLLQAEIEELRATLEQTERSRKIAEQELLDASERVQLLHTQNTS    1726
sp|P02567|MYO1_CAEEL      RREEIRENYLAAEKRLAIALSESEDLAHRIEASDKHKKQLEIEQAELKSSNTELIGNNAA    1729
                           :*:::*:   .*:*  :  :* *:*   :* ::: :*  * *  : ..    *  :*::

sp|P12882|MYH1_HUMAN      LINTKKKLETDISQIQGEMEDIIQEARNAEEKAKKAITDAAMMAEELKKEQDTSAHLERM    1786
sp|P02567|MYO1_CAEEL      LSAMKRKVENEVQIARNELDEYLNELKASEERARKAAADADRLAEEVRQEQEHAVHVDRQ    1789
                          *   *:*:*.::.  :.*::: ::* : :**:*:** :**  :***::**: :.*::*

sp|P12882|MYH1_HUMAN      KKNLEQTVKDLQHRLDEAEQLALKGGKKQIQKLEARVRELEGEVESEQKRNVEAVKGLRK    1846
sp|P02567|MYO1_CAEEL      RKSLELNAKELQAKIDDAERAMIQFGAKALAKVEDRVRSLEAELHSEQRRHQESIKGYTK    1849
```

```
                          :*.** ..*:** ::*:**:  :: * * : *:* ***.**.*:.***:*: *::**  *

sp|P12882|MYH1_HUMAN      HERKVKELTYQTEEDRKNILRLQDLVDKLQAKVKSYKRQAEEAEEQSNVNLSKFRRIQHE    1906
sp|P02567|MYO1_CAEEL      QERRARELQFQVEEDKKAFDRLQENVEKLQQKIRVQKRQIEEAEEVATQNLSKFRQIQLA    1909
                          :**:.:** :*.***:* : ***: *:*** *::  *** ***** :. ******:**

sp|P12882|MYH1_HUMAN      LEEAEERADIAESQVNKLRVKSREVHTKIISEE 1939
sp|P02567|MYO1_CAEEL      LENAEERAEVAENSLVRMRGQVVRSATNK---- 1938
                          **:*****::**..: ::* : . *:
```

**Figure S3: Conservation of MYO-1 methylation sites in human myosin-1.**

A Clustal Omega (v1.2.4) multiple sequence alignment was used to align *C. elegans* MYO-1 (MYO1_CAEEL) with the human ortholog myosin-1 (MYH1_HUMAN) to assess conservation of methylation sites detected in our dataset. Methylated residues identified in MYO-1 were mapped onto the alignment and compared with the corresponding positions in human myosin-1. The following MYO-1 lysine sites align to conserved lysines in MYH1: K71→K73, K188→K190, K642→K639, K645→K642, and K1544→K1541, whereas K55 and K1539 did not map to a corresponding lysine in MYH1.

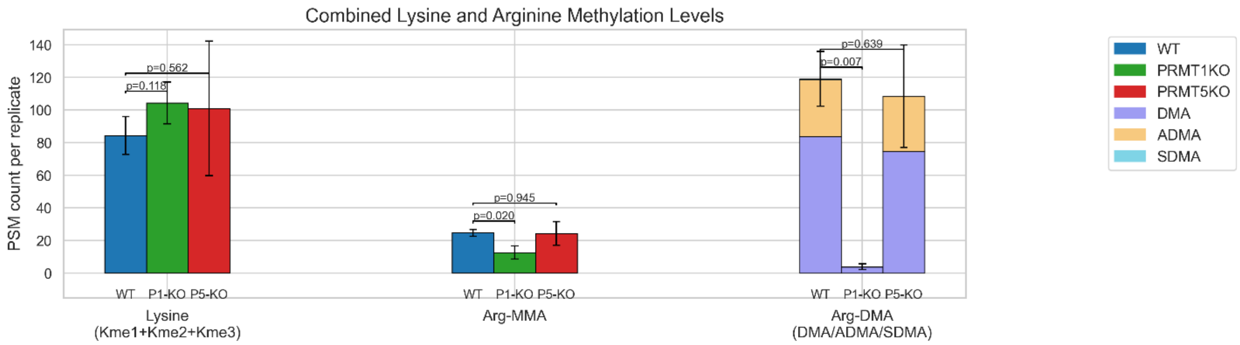


**Figure S4: PSM counts of lysine and arginine methylation.**

PRMT-1 knockout selectively reduces protein arginine methylation while lysine methylation remains unchanged. Bar plots show the number of methylated peptide-spectrum matches (methyl-PSMs) passing the FDR threshold for lysine methylation (Kme1, Kme2, and Kme3) and arginine methylation across WT, *prmt-1*-KO, and *prmt-5*-KO samples. Total lysine methyl-PSM counts were comparable among all three genotypes. In contrast, arginine monomethylation (MMA) was strongly reduced in *prmt-1*-KO, and arginine dimethylation (DMA, assigned as ADMA and SDMA where possible based on neutral loss) was nearly undetectable in *prmt-1*-KO. No statistically significant differences were observed for *prmt-5*-KO relative to WT. Bars represent mean ± SD across biological replicates; p-values are indicated for the comparisons shown.

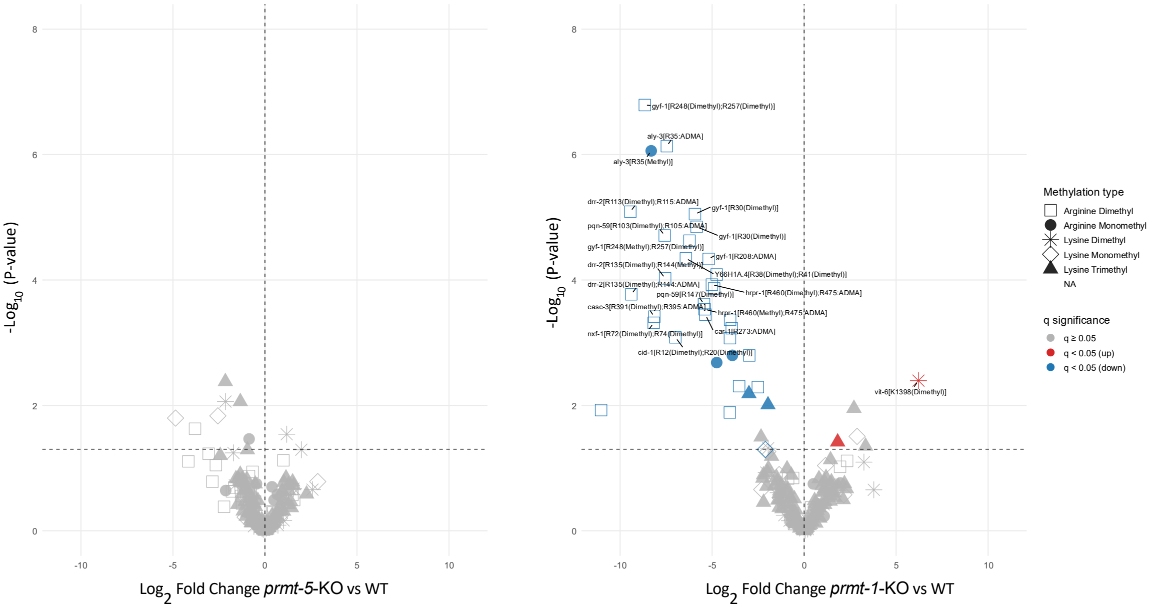


**Figure S5: Differential methyl-peptide abundance in *prmt-5*-KO versus WT and *prmt-1*-KO versus WT.**

(A) Volcano plot of quantified methyl-peptides comparing *prmt-5*-KO to WT. No methyl-peptides met the 5% FDR threshold ($q < 0.05$), consistent with the lack of separation between WT and *prmt-5*-KO observed by PCA. (B) Volcano plot of quantified methyl-peptides (both methyl-arginine and methyl-lysine) comparing *prmt-1*-KO to WT. A subset of methyl-peptides showed significant changes at $q < 0.05$ (blue, decreased in *prmt-1*-KO; red, increased in *prmt-1*-KO).

```
sp|O43390|HNRPR_HUMAN      ------------------------------MANQVNGNAVQLKEEEEPMDTSSVTHTEHY  30
tr|Q9NLD1|Q9NLD1_CAEEL     MSEEIKQEVKEEAPETTTEAKTEEAAPVAEVVAEVVTNGDKPAENGKTEEPSKYAETENY  60
                                                          :. :*  *. :  *: :  : *. :.**:*

sp|O43390|HNRPR_HUMAN      KTLIEAGLPQKVAERLDEIFQTGLVAYVDLDERAIDALREFNEEGALSVLQQFKESDLSH  90
tr|Q9NLD1|Q9NLD1_CAEEL     KKLIGLKMKPALADVVCEIYESGLLGPEDLDDRAVDIINSVNLDQAKFIFTEIKNSELFG  120
                           *.**   :   :*: : **:::**:.  ***:**:* :...* : *  :: ::*:*:*

sp|O43390|HNRPR_HUMAN      VQNKSAFLCGVMKTYRQREKQGS----KVQESTKGPDEAKIKALLERTGYTLDVTTGQRK  146
tr|Q9NLD1|Q9NLD1_CAEEL     VATKSLYVTSLIRSFKDRCRQQGAAAVTSGKLINGPELAALKNLLETTGYTIEVTIGQRK  180
                           * .** :: .:::::::* :* .    .  :  :**: * :* *** ****::** ****

sp|O43390|HNRPR_HUMAN      YGGPPPDSVYSGV-QPGIGTEVFVGKIPRDLYEDELVPLFEKAGPIWDLRLMMDPLSGQN  205
tr|Q9NLD1|Q9NLD1_CAEEL     FGGPPPDWEGPATGPAGQGHEIYVGHIPTDVFEDTLVPLFEKSGKIWDLRLMMDPMSGAS  240
                           :******    ..   * * *::**:** *::** *******:* **********:** .

sp|O43390|HNRPR_HUMAN      RGYAFITFCGKEAAQEAVKLCDSYEIRPGKHLGVCISVANNRLFVGSIPKNKTKENILEE  265
tr|Q9NLD1|Q9NLD1_CAEEL     RGYAFVTYCNKEDAAAAAKTYDGHEISTGKPLKVNVSIANTRLFIGNIPKTKSKDEILEE  300
                           *****:*:*.** *  *.*  *.:**  ** * * :*:**.***:*.***.*:*::****

sp|O43390|HNRPR_HUMAN      FSKVTEGLVDVILYHQPDD-KKKNRGFCFLEYEDHKSAAQARRRLMSGKVKVWGNVVTVE  324
tr|Q9NLD1|Q9NLD1_CAEEL     LKTHAEGVVDVIVYSVPDNEKIKNRGFCFVDFIDHKTASDIKRKIAQHKIRPFNADVYVD  360
                           :.. :**:****:*  **: * *******::: ***:*:: :*:: . *:: :.  * *:

sp|O43390|HNRPR_HUMAN      WADPVEEPDPEVMAKVKVLFVRNLATTVTEEILEKSFSEFGKLERVKKLKDYAFVHFEDR  384
tr|Q9NLD1|Q9NLD1_CAEEL     WAEHQEEPDEDTMSKVKVLYIRNIKEAVTEEKLNELFKEYASLDRVKKVKDYAFIHFNER  420
                           **:  **** :.*:*****::**:  :**** *:: *.*:..*:****:*****:**::*

sp|O43390|HNRPR_HUMAN      GAAVKAMDEMNGKEIEGEEIEIVLAKPPDKKRKERQAARQASRSTAYEDYYYHPPPRMPP  444
tr|Q9NLD1|Q9NLD1_CAEEL     DDCLKAMEEWNGKELEGTVVEASLAKPPQEKKKKPV------------------------  456
                           . .:***:* ****:**  :*  *****::*:*:

sp|O43390|HNRPR_HUMAN      PIRGRGRGGGRGGYGYPP------------DYYGYEDYYDDYYG--YDYH-DYRGGYEDP  489
tr|Q9NLD1|Q9NLD1_CAEEL     -MRGRGFGGAGGGAGNHGQRGGQRGGGQWGGNAGPGGYYPNHFNNGYDMPMPWGGGY---  512
                            :**** **. ** *               .  *  .** :::.  **    : ***

sp|O43390|HNRPR_HUMAN      YYGYDDGYAVRG--RGGGRGGRGAPPPPRGRGAPPPRGRAGYSQRGAPLGPPRGSRGGRG  547
tr|Q9NLD1|Q9NLD1_CAEEL     --GGDMGYGAYGGGYGGGYGGPGAYGDF---G---AYG--GGGFRGGMRGSPRGGMGGGG  562
                             * * **.. *   *** ** **       *     *  * . **.  * ***. ** *

sp|O43390|HNRPR_HUMAN      GPAQQQRGRGSRGSRGNRGGNVGGKRKADGYNQPDSKRRQTNNQNWGSQPIAQQPLQQG  607
tr|Q9NLD1|Q9NLD1_CAEEL     GFRGGPPGRGGMANRRGR-----GKRPGDGRGGPASKRDNGK--PDFSAD-VNMSTF---  611
                           *      ***. ..* .*     *** .** . * *** : :   ::.:: :  . :

sp|O43390|HNRPR_HUMAN      GDYSGNYGYNNDNQEFYQDTYGQQWK      633
tr|Q9NLD1|Q9NLD1_CAEEL     --------------------------      611
```

**Figure S6: HRPR-1(Q9NLD1)/hnRNPR (HNRPR) alignment highlights PRMT1-targetable RGG/RG motifs.**

A Clustal Omega (v1.2.4) multiple sequence alignment of *C. elegans* HRPR-1 and human hnRNPR is shown. Putative RGG/RG motif regions are highlighted (human, red; worm, blue), and bolded arginines denote quantified sites that exhibit reduced methylation in *prmt-1*-KO worms.

```
sp|P35637|FUS_HUMAN         MASNDYTQQATQSYGAYPTQPGQGYSQQSSQPYGQQSYSGYSQSTDTSGYGQSSYSSYGQ  60
tr|Q18265|Q18265_CAEEL      ------------------------------------------------------------  0

sp|P35637|FUS_HUMAN         SQNTGYGTQSTPQGYGSTGGYGSSQSSQSS-YGQQSSYPGYGQQPAPSSTSGSYGSSSQS  119
tr|Q18265|Q18265_CAEEL      -----------------MAAYDQSQPDYSTPEGQQAYWAYY-------------QQQQQQ  30
                                              ..*..** . *:  ***: :  *              ...*.

sp|P35637|FUS_HUMAN         SSYGQPQSGSYSQQPSYGGQQQSYGQQQSYNPPQGYGQQNQYNSSSGGGGGGGGGGNYGQ  179
tr|Q18265|Q18265_CAEEL      QPGGQPDQDPYA-AAAYGGHDQAQQPQNPYAPPPPGA--DPYGQGSG--GQSGGSDPYGQ  85
                            .  ***:.. *:   :***::*:   *: * **   .  : *...**  * .**.. ***

sp|P35637|FUS_HUMAN         DQSSMSSGGGSGGGYGNQDQ------SGGGGSGGYGQQDRGGRGRGGSGGGGGGGGGGYN  233
tr|Q18265|Q18265_CAEEL      SR-----GGG-RGGFGGSRGGGGYDGGRGGSRGGYD---GGRGGYGGDRGGRGGGRGGYD  136
                            .:     ***  **:*..        . **. ***.    *  * **. ** *** ***:

sp|P35637|FUS_HUMAN         RSSGG---YEP---RGRGGGRGGRGGMGG--------SDRGGFNKFGGPRDQGSRHDSEQ  279
tr|Q18265|Q18265_CAEEL      GERRGGSRWDDGNSDRQGGPPGGRGGYQDRGPRRDGPPSGGGYGG-GGAASGNREFGSDG  195
                             .  *   ::      :**  *****  .         . **:.  **  . . ...*:

sp|P35637|FUS_HUMAN         DNSDNNTIFVQGLGENVTIESVADYFKQIGIIKTNKKTGQPMINLYTDRETGKLKGEATV  339
tr|Q18265|Q18265_CAEEL      RVELKETVFVQGISTTANEAYIADVFSTCGDIAKN--DRGPRIKIYTDRNTGEPKGECMI  253
                              . ::*:****:. ...   :** *.  * * .*     * *::****:**: ***. :

sp|P35637|FUS_HUMAN         SFDDPPSAKAAIDWFDGKEFSGNPIKVSF--ATRRADFN--RGGGNGR--GGRGRGGPMG  393
tr|Q18265|Q18265_CAEEL      TFVDASAAQQAITMYNGQPFPGGSSPMSISLAKFRADAGGERGGRGGRGGFGGGRGGPMG  313
                            :* *  :*: **  ::*: * *.   :*:  *. *** .  *** .**   * *******

sp|P35637|FUS_HUMAN         R--------GGYGGGGSG------------------------------------------  403
tr|Q18265|Q18265_CAEEL      GRGGFGGDRGGYGGGGGRGGFDGGRGGGGGFRGGDRGGFRGGDRGGFRGGDRGGFRGGDR  373
                                     *******.

sp|P35637|FUS_HUMAN         GGGRGGFPSG----GGGGGGQQRAGDWKCPNPTCENMNFSWRNECNQCKAPKPDGPGGGP  459
tr|Q18265|Q18265_CAEEL      GGDRGGFRGGRGVGGGNANMEQRKNDWPCE--QCGNSNFAFRRECNQCQAPRPDGGSGGG  431
                            **.**** .*    **... :** .** *    * * **::*.*****:**:*** .**

sp|P35637|FUS_HUMAN         GGSHMGGNYGDDRRGGRGGYDRGGYRGRGGDRGGFRGGRGGGDRGGFGPGKMDSRGEHRQ  519
tr|Q18265|Q18265_CAEEL      GGERRGGPPGGDRYRPY-------------------------------------------  448
                            **.: **  *.**

sp|P35637|FUS_HUMAN         DRRERPY   526
tr|Q18265|Q18265_CAEEL      -------   448
```

**Figure S7: FUST-1/FUS alignment places the PRMT-1-dependent FUST-1 R160 site within the conserved RGG1 motif region of human FUS.**

A Clustal Omega (v1.2.4) multiple sequence alignment of *C. elegans* FUST-1 and human FUS is shown. The RGG1 motif of human FUS is highlighted in red, and FUST-1 R160 (ADMA site; reduced in *prmt-1*-KO) is indicated in bold blue. Alternative alignment approaches can map R160 to nearby positions within the same RGG1 region (e.g., R244 using the ExPASy SIM tool with the BLOSUM62 matrix); the SIM alignment is not shown here.

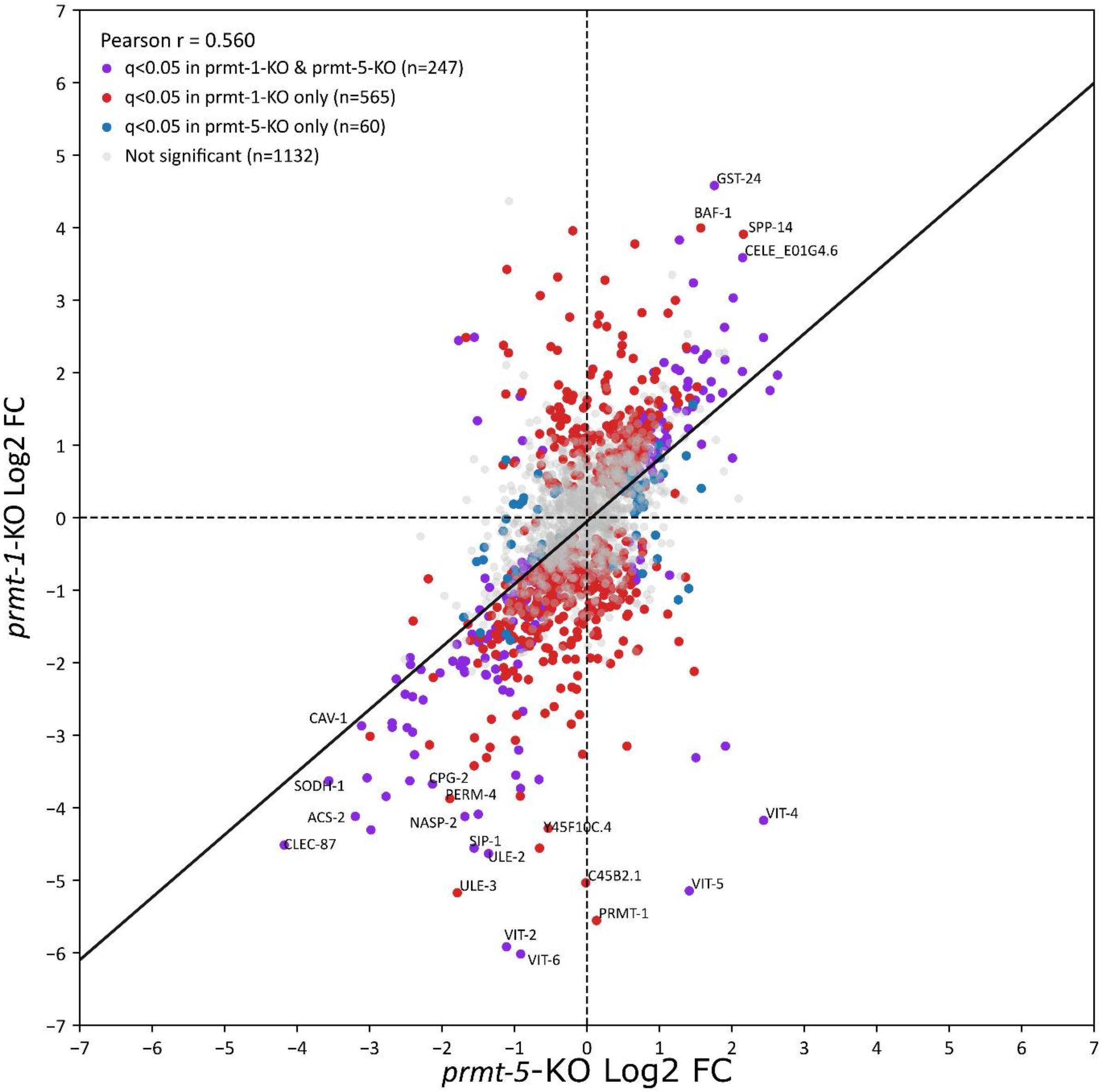


**Figure S8: Concordant protein-level changes in *prmt-1*-KO and *prmt-5*-KO worms.**

Scatter plot comparing $\log_2$ fold-changes (log2FC) for proteins quantified in both datasets (x-axis: *prmt-5*-KO vs WT; y-axis: *prmt-1*-KO vs WT). Points are colored by significance (q < 0.05) in *prmt-1*-KO and/or *prmt-5*-KO, with non-significant proteins shown in gray. A linear fit is overlaid, highlighting that proteins significantly regulated in both genotypes tend to change in the same direction (Pearson r = 0.56). Selected proteins with the largest deviations are annotated.
